# A citizen science photo guide of plants and bee visitors in the Eastern Afromontane Biodiversity Hotspot of Kenya

**DOI:** 10.64898/2025.12.04.692413

**Authors:** Fairo F. Dzekashu, Marcell K. Peters, Ingolf Steffan-Dewenter, Jayne M. Macharia, Kennedy W. Matheka, Abdullahi A. Yusuf, Christian W.W. Pirk, H. Michael G. Lattorff

## Abstract

The influence of global change on species assemblage patterns is on the rise with evidence showing a steady decline in pollinator communities across the globe. As such, the implementation of rigorous monitoring efforts aimed at documenting the actual status of pollinators is of essence especially in the tropics. This can be possible through participatory studies with volunteers (citizen scientists) that emphasizes on apriori research hypotheses with the aim to draw inferences on the potential drivers of species community assemblage patterns.

In ecology, photos have the ability to drive change and can provide unbiased evidence on species existence with a potential to compare changes over time and space. However, in the tropics and particularly in sub-Saharan Africa, such ideas fall short of public participation. In this study, we aimed to assess and digitally document plants and their bee visitors as important data sources with immense potential to improve on the knowledge and awareness of the different plants and bee species in two counties of Kenya (Taita and Murang’a). Our results showed marked differences in bee community assemblage patterns between the two counties, with a huge number of unique species between them. We successfully digitalised and present photos of 114 bee and 262 plant species. Using these digitalised photographs, we herein elaborate on the community assemblage patterns of plants and their bee pollinators in the Eastern Afromontane Biodiversity Hotspot (EABH) of Kenya and further discuss the importance of exploiting images for research in biodiversity and conservation. We therefore conclude that, technological advances that helps in digitalising species through photography can be important in the successful implementation of community based environmental threat management agendas.

## Introduction

Insect pollinators are important contributors in the fertilization and reproduction of crop and wild plants, thus contributing immensely to ecosystem stability and ensuring food security (Ollerton et al., 2011). However, compelling evidence continue to reveal a steady decline in pollinator communities across the world. These declines can be attributed to interrelated anthropogenic activities such as agricultural intensification, indiscriminate use of pesticides, loss of natural vegetations due to the clearing of natural vegetations, the introduction of invasive alien species and pathogens, and climate change (Powney et al., 2019; Potts et al., 2016; Vanbergen et al., 2013). In the face of the ever-increasing global demand for insect pollinated crops and wild plants, a resounding decline in pollinator species would have far reaching economic and health related consequences because of shortages in crops and medicinal plants (Eilers et al., 2011). Majority of countries in the northern hemisphere have in place rigorous and efficient strategies to monitor and document the status of plants and pollinator species (Lanuza et al., 2025; Rondeau et al., 2023; Woodard et al., 2020), with many of the plants and their pollinators well known and documented. On the other hand, there is a huge absence of historical and contemporary data of plants and insect pollinators that can be used to compare community changes or shifts in distributions across the African continent as only a few recent efforts have been aimed at documenting the occurrences of plants and their insect pollinators across the different gradients of the continent (Dzekashu et al., 2022; Guy et al., 2021; Mertens et al., 2021; Stein et al., 2021; Lasway et al., 2022; Baldock et al., 2011).

Amidst alarming concerns around species declines, diversity hotspots like the Eastern Afromontane Biodiversity Hotspot (EABH) are important refugia for species (Dzekashu et al., 2022). It is mostly espoused that the EABH harbours considerable amount of plant and pollinator species that extends across vast spatial terrains represented by natural vegetations and anthropogenically induced landscapes, from the Aberdare mountains to the Taita hills. Sharp changes in climate, vegetation, anthropogenic and cultural activities along the slopes of the EABH encourages the estimation of species community assemblage patterns at short spatial scales. Moreover, quantifying the number and occurrence of species in this region is of critical interest as such knowledge would contribute to the understanding of the eco-evolutionary processes driving species persistence (as some would increase in relative abundance), and the crucial ecosystem services they provide. This knowledge would also guide on the appropriate implementation of policies and management plans geared at conserving declining species. Despite recent studies on plant-pollinator community assemblage patterns in this region, much remains to be explored (Baldock et al., 2011; Guy et al.,2021; Seifert et al., 2022; Dzekashu et al.,2023), as currently, it is not well understood how the composition of species communities varies along the different slopes of the EABH. Assembling the public through participatory dialogues with the aim of developing more engaging and well-grounded decision-making programs is important in the successful implementation of community based environmental threat management agendas (Bonney et al., 2009; Alan Irwin, 1995). Such assembles are best in a synergy between professional research scientists and members of the public when collectively they implement a unified strategy with the aim of collecting and processing data that can be useful in designing public policies (Bonney et al., 2014). Under such synergies, professional scientists would train public participants or volunteers on standard research protocols and ethics aimed at addressing specific research questions, methods or designs required for the study (e.g. selection of sampling plot(s) and the reason for selecting such plot(s), and importantly, ensure the reproducibility of any sampling techniques used), data collection, processing and interpretation. Such procedures can be very useful for long-term biodiversity monitoring studies in the tropics, where there exist huge deficiencies in public understanding of experimental protocols, which also contributes in reducing enthusiasm for participations in research-based studies with potentials of improving the well-being of individuals and the local communities. Therefore, citizen science studies on biodiversity in the tropics can be designed with apriori hypotheses such that they not only emphasise on the status of priority species but also emphasises on assessing and identifying species community assemblage patterns and trends, drawing inferences on the potential drivers of such patterns (Dickinson et al., 2010).

Images are capable of driving rapid actions and can influence positive changes, particularly in the current plight met by endangered species. In ecology, photos can serve as compelling proof of the effects of global change on biodiversity (Depauw et al., 2022). For species, image data can be more time and cost effective, and may provide novel insights that are not possible with other data formats (Gonella et al., 2015). They can also provide unbiased evidence on species existence with a potential to compare changes over time and space (Sanseverino et al., 2016). Additionally, they can serve as useful learning and referenced sources for citizen scientists as they try to classify sampled organisms. For professional scientists, species images (especially those with georeferenced information) could facilitate mapping of species occurrences and thereby enabling the prediction of their spatio-temporal distributions (ElQadi et al., 2017).

During the design and sampling events for our study on plant-pollinator interaction networks on the Afromontane Biodiversity Hotspots in Kenya (Dzekashu et al., 2022, 2023, 2024, 2025), we worked with several local residence and remarked a high level of unawareness of bee pollinators other than honeybees. We here aimed to understand and document plants and their bee visitors along two mountain slopes (i.e. Taita Hills and Murang’a) of the Eastern Afromontane Biodiversity Hotspot (EABH) in Kenya. To achieve this, we collected and present pictographic evidence of plants and their bee visitors as important data sources with immense potential to improve on the knowledge and awareness of the different plants and bee species in the local communities of these two regions of Kenya and to highlight their conservation status.

## Methods

We conducted sampling along two elevation gradients situated within the Eastern Afromontane Biodiversity Hotspot (EABH) in Kenya; the Taita Hills located in Taita Taveta county (38°10‘ to 39°03’E,-3°15‘ to-4°0’S, 525-1,865 m above sea level) and in Murang’a County, located in the central region of Kenya (0°34‘ to 1°5’S, 36°43‘ to 37°27’E, 1, 470-2,530 m above sea level) (Figure 1 a-a2). This area consists of a sub-tropical climate with arid and semi-arid conditions and savannah-like vegetation at lower elevations whilst montane and premontane forests occupy higher elevations. The mean annual minimum and mean annual maximum temperatures are ∼17.5 to 19 °C and ∼29 to 31°C (Gebrechorkos et al., 2019), while the mean annual precipitations are ∼250 mm to ca. 2,000 mm (Orodho, 2006) from low to high elevations. Intensive sampling was conducted in 50 selected study plots of 100 m × 100 m size, that were equally distributed among the two elevation gradients. All plots were positioned within regrowth vegetation with elevational increments of ca. 100-250 m among next neighbouring plots. The minimum linear distance between adjacent plots was ca. 2.3 km.

**Figure 1.**
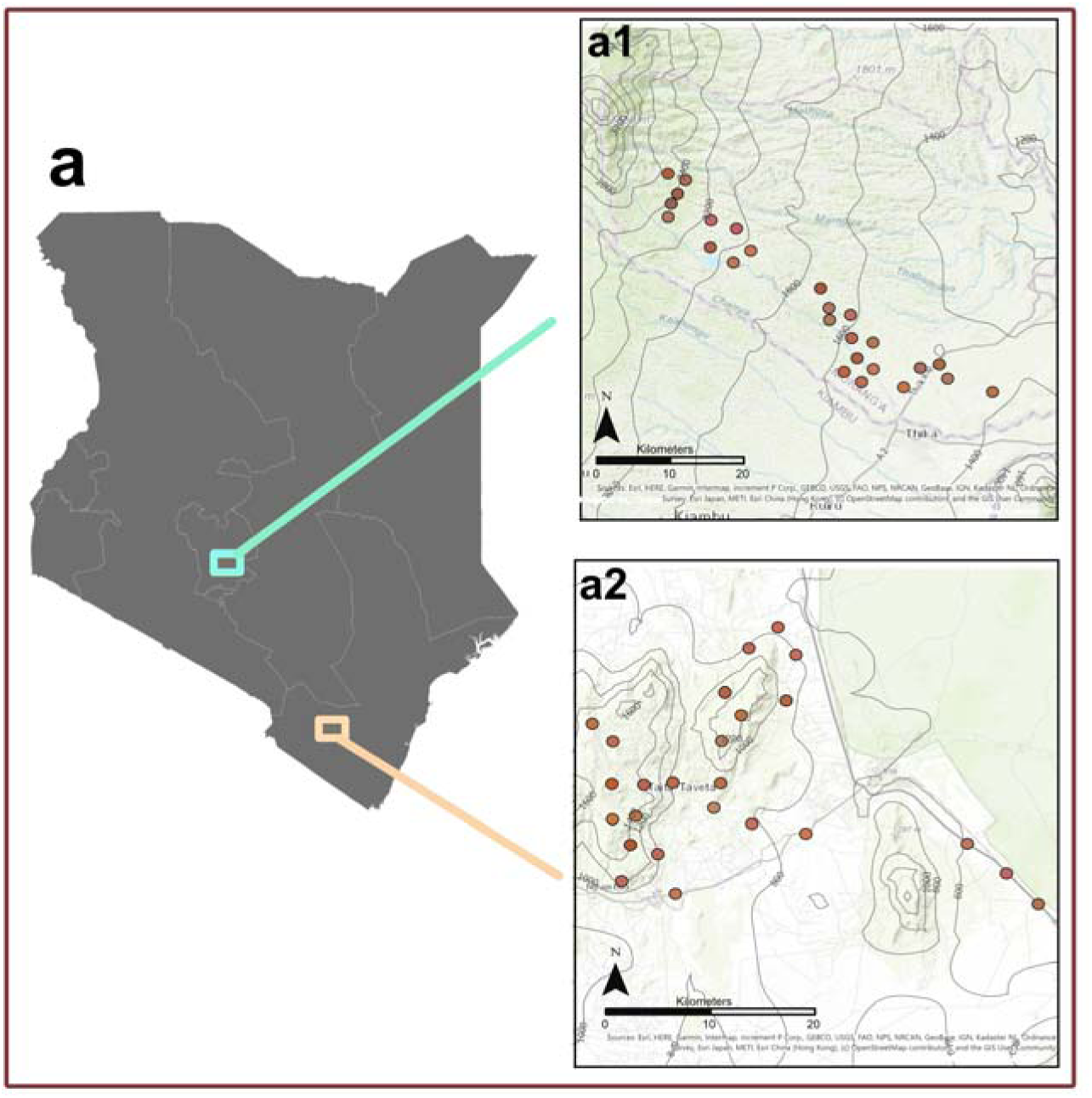
Represents the map of Kenya and the two studied counties (a1: Murang’a & a2: Taita) and brown dots within counties represents sampling plots

**Figure 2.**
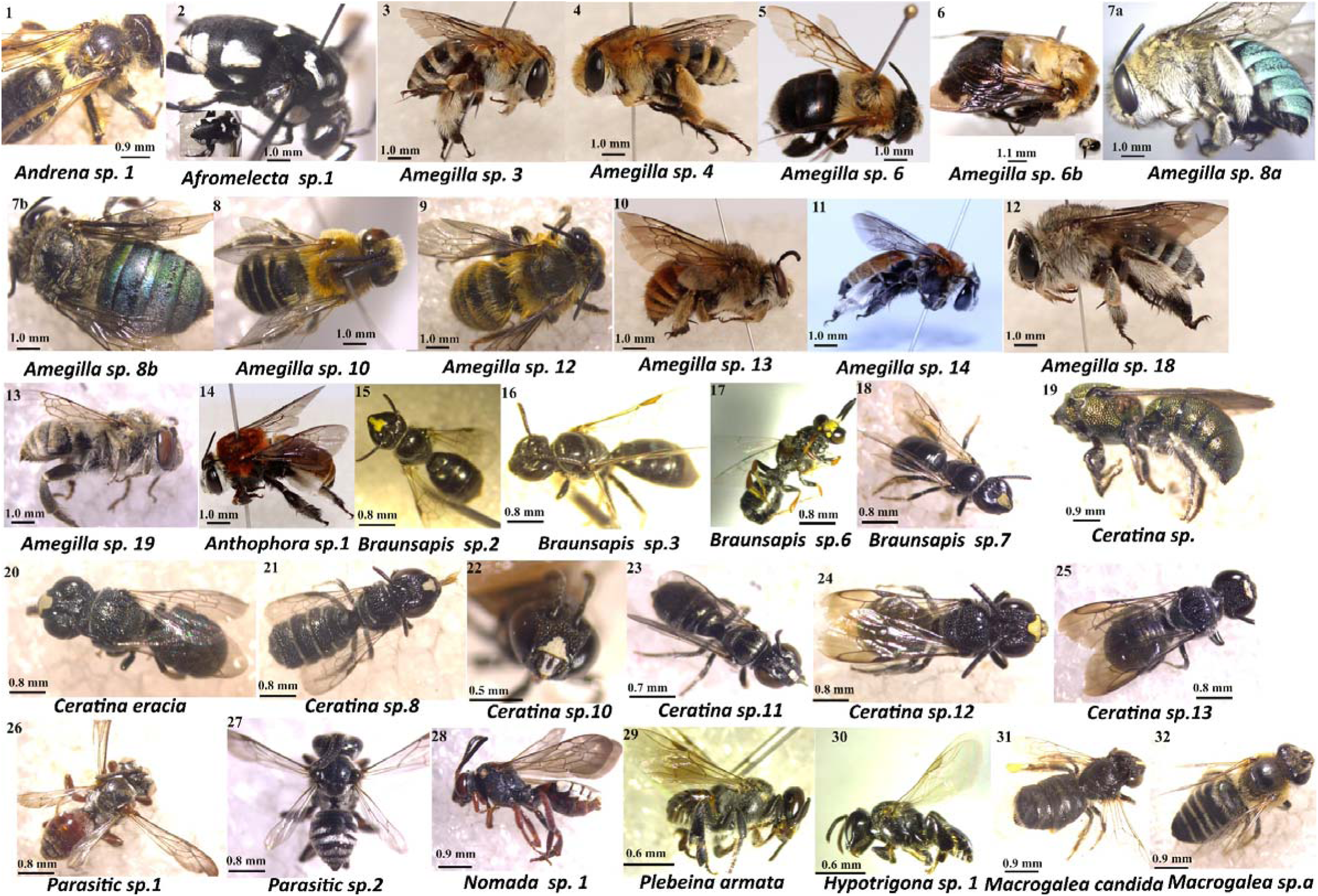
Bee species found visiting plants on study plots. Bee species are arranged according to families; 1: Andrenidae, 2-32: Apidae. Measurement in mm indicates the scale of each photo. Photo credit: Fairo Dzekashu

**Figure 3.**
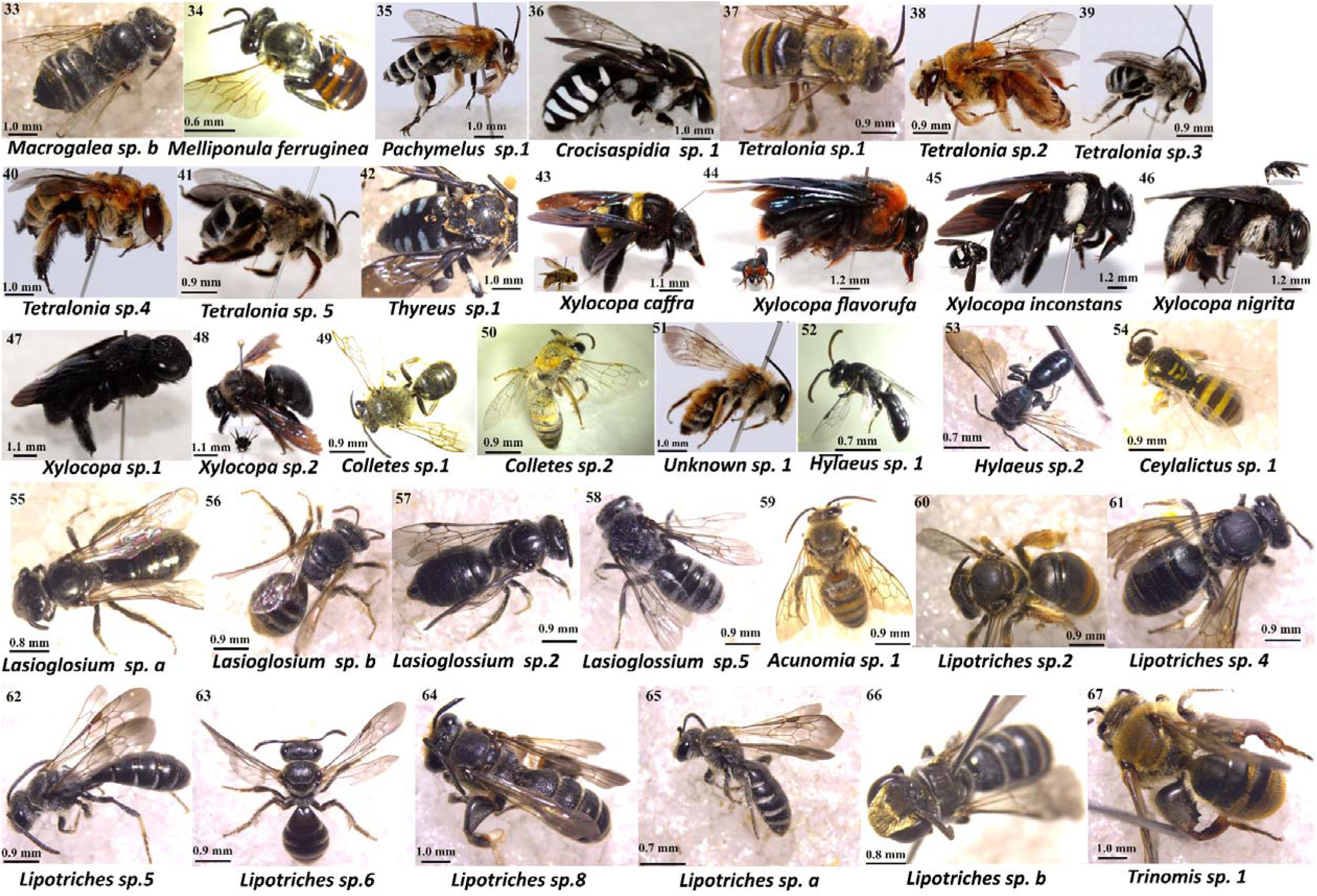
Bee species found visiting plants on study plots. Bee species are arranged according to families; 33-48: Apidae, 49-53: Colletidae, 54-67: Halictidae. Measurement in mm shows the scale of each photo. Photo credit: Fairo Dzekashu

**Figure 4.**
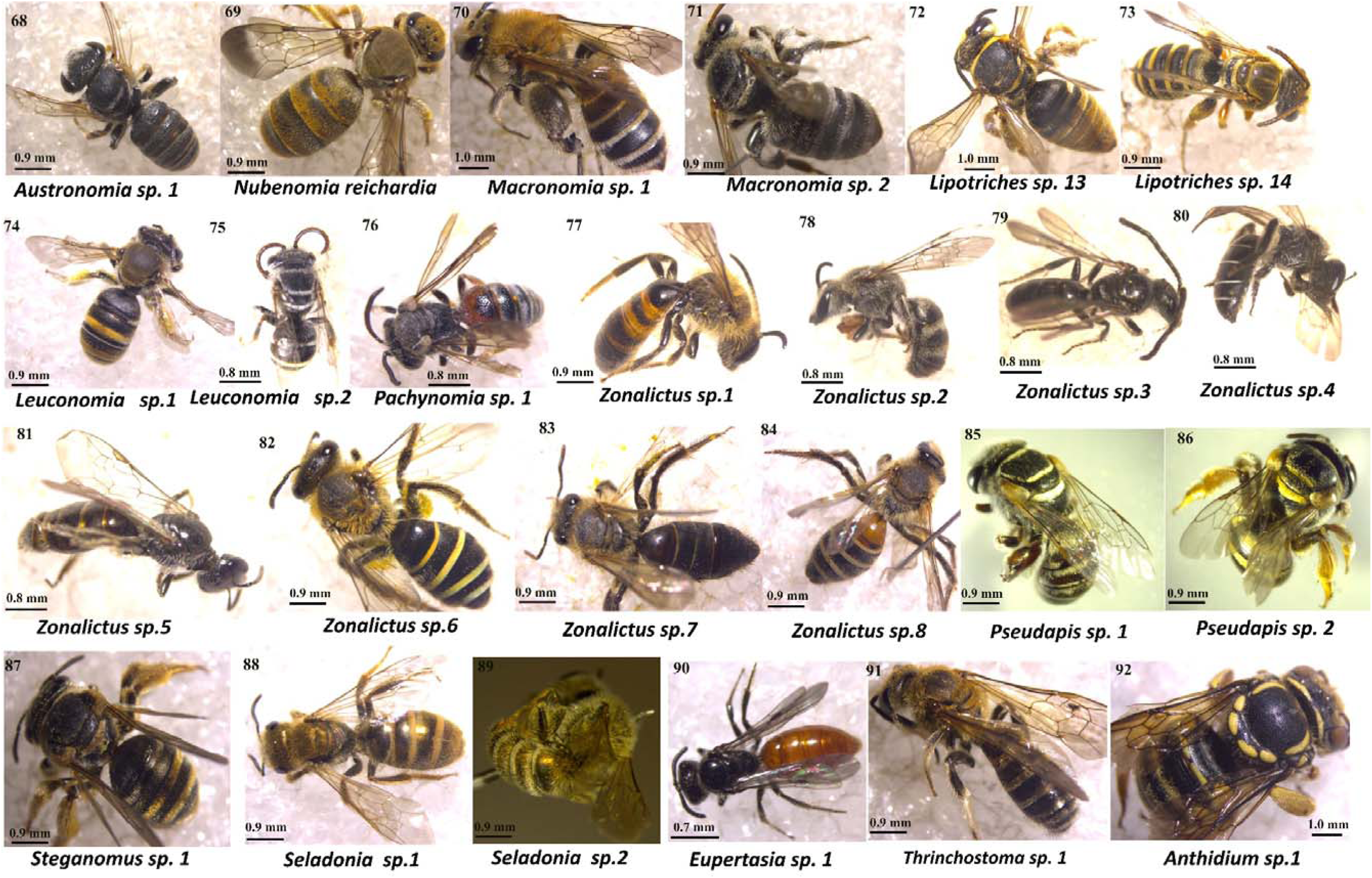
Bee species found visiting plants on study plots. Bee species are arranged according to families; 68-91: Halictidae, 92: Megachilidae. Measurement in mm indicates the scale of each photo. Photo credit: Fairo Dzekashu

**Figure 5.**
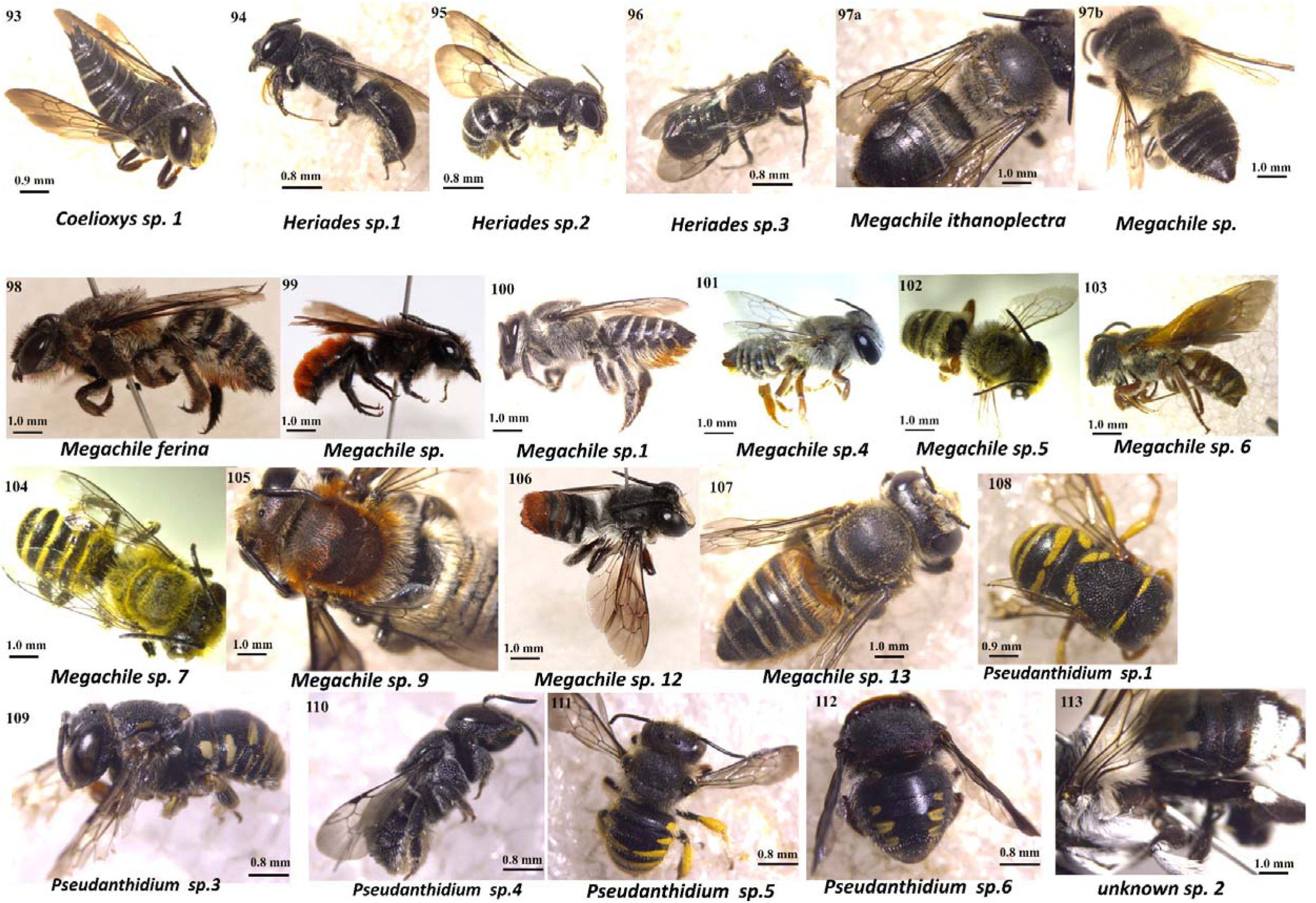
Bee species found visiting plants on study plots. Bee species are arranged according to families; 93-113: Megachilidae. Measurement in mm shows the scale of each photo. Photo credit: Fairo Dzekashu

**Figure 6.**
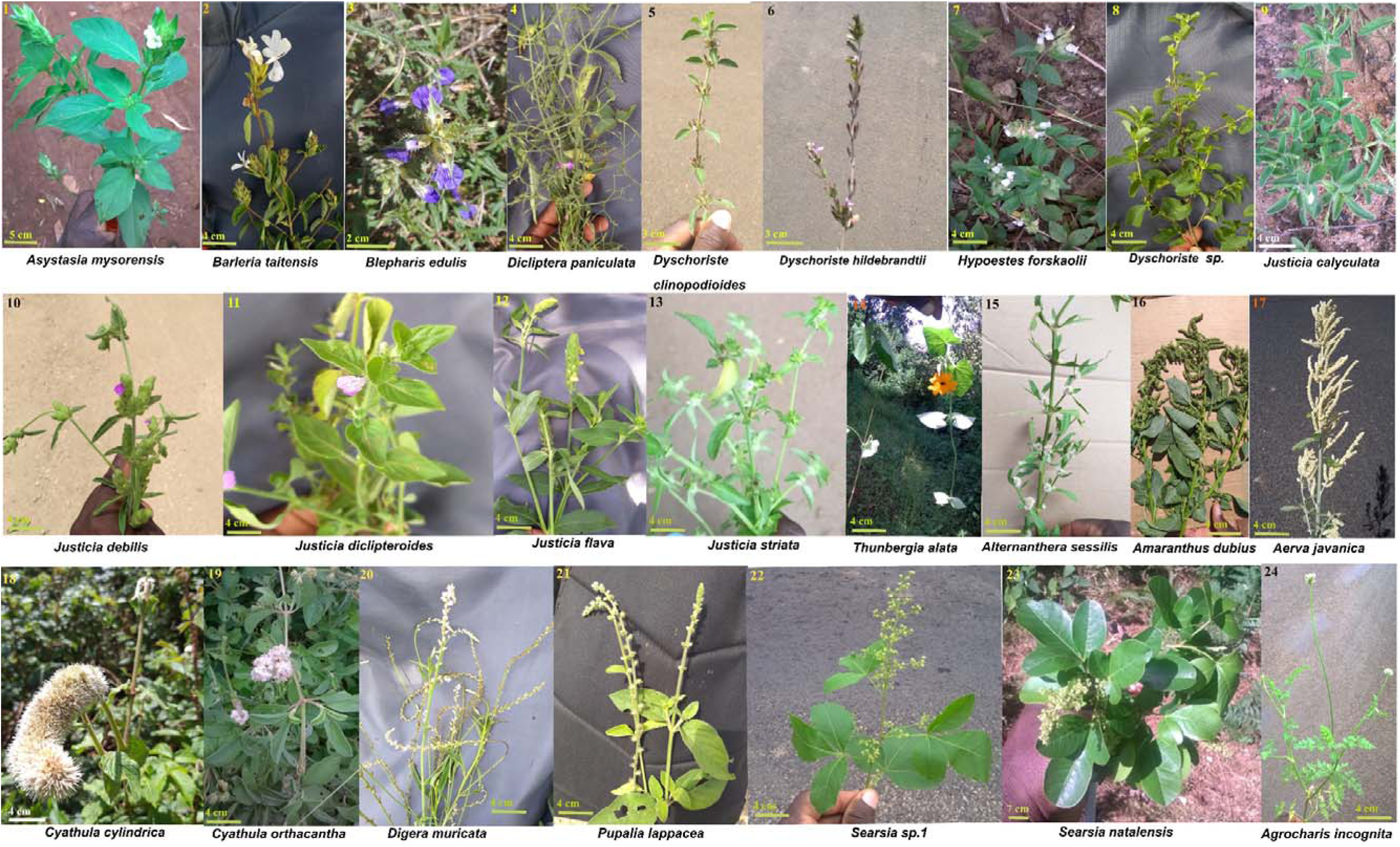
Plant species on which bee species were found visiting on study plots. Plant species are arranged according to families; 1-14: Acanthaceae, 15 – 23: Amaranthaceae, 24: Apiaceae. Measurement in cm depicts the scale of each photo. Photo credit: Fairo Dzekashu.

**Figure 7.**
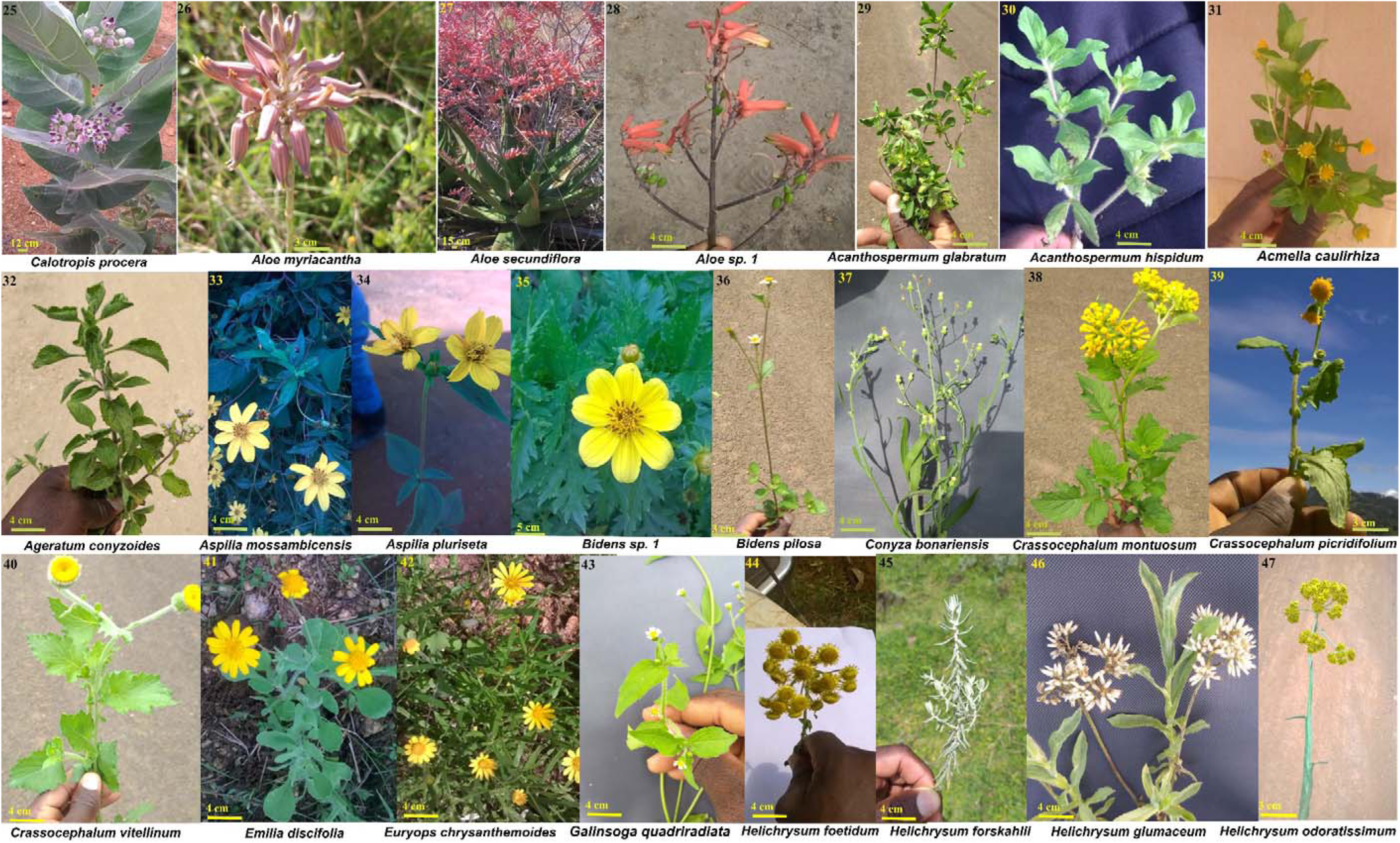
Plant species on which bee species were found visiting on study plots. Plant species are arranged according to families; 25: Apocynaceae, 26-28: Asphodelaceae, 29-47: Asteraceae. Measurement in cm shows the scale of each photo. Photo credit: Fairo Dzekashu

**Figure 8.**
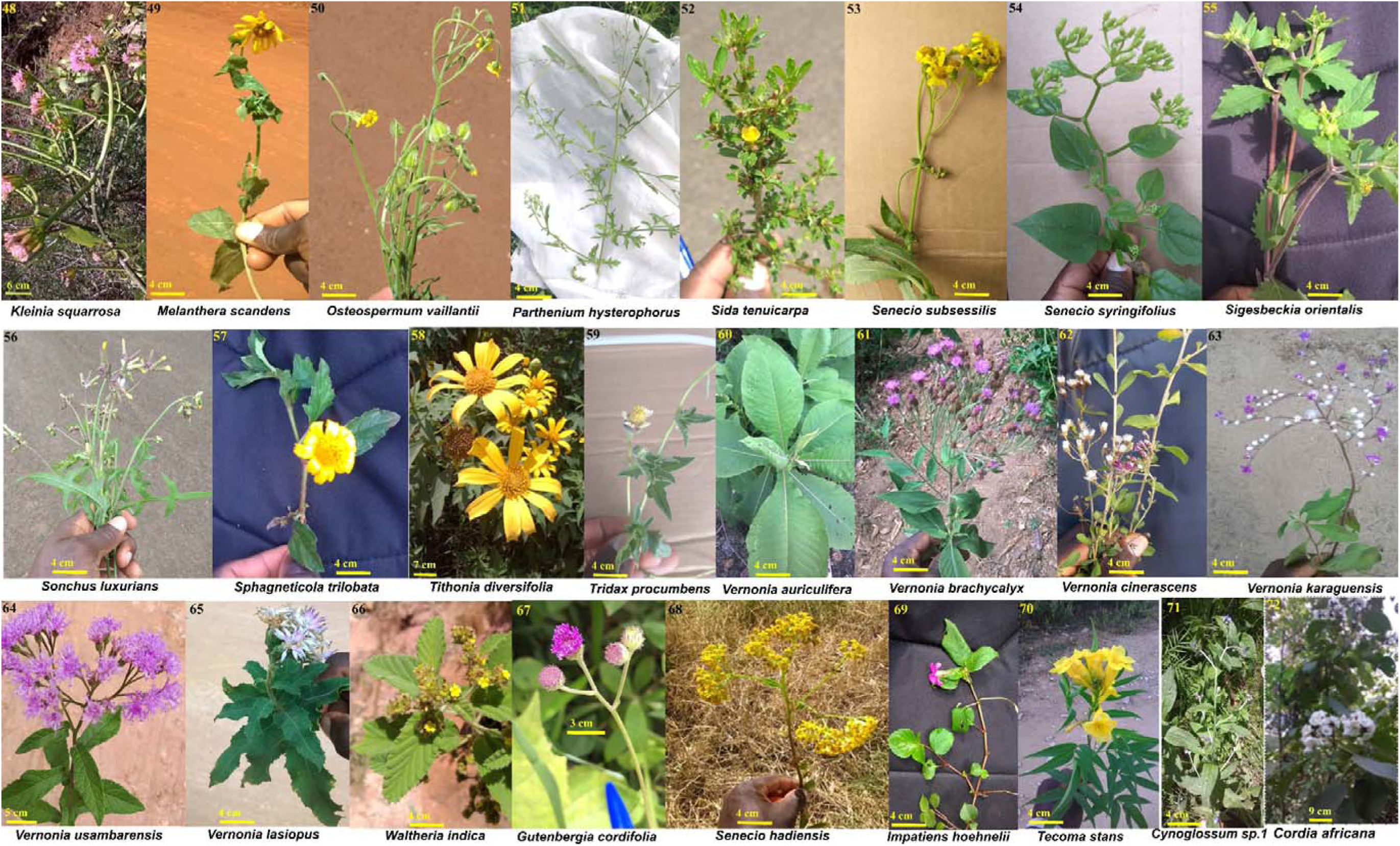
Plant species on which bee species were found visiting on study plots. Plant species are arranged according to families; 48-51 & 53-68: Asteraceae, 69: Balsaminaceae, 70: Bignoniaceae, 71-72: Boraginaceae, Measurement in cm indicates the scale of each photo. Photo credit: Fairo Dzekashu

**Figure 9.**
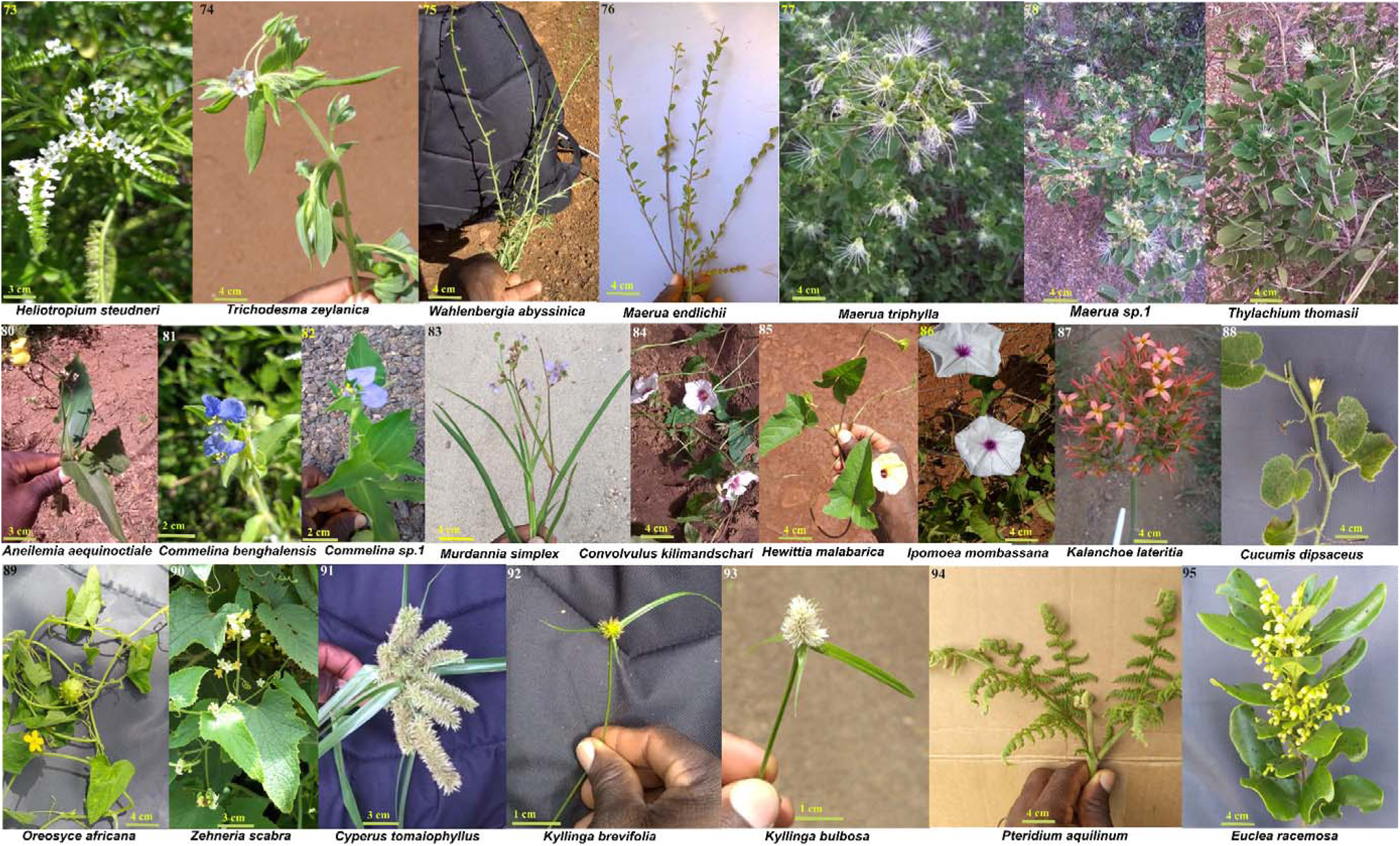
Plant species on which bee species were found visiting on study plots. Plant species are arranged according to families; 73-74: Boraginaceae, 75: Campanulaceae, 76-79: Capparaceae, 80-83: Commelinaceae, 84-86: Convolvulaceae, 87: Crassulaceae, 88-90: Cucurbitaceae, 91-93: Cyperaceae, 94: Dennstaedtiaceae, 95: Ebenaceae. Measurement in cm depicts the scale of each photo. Photo credit: Fairo Dzekashu.

### Bees and plants sampling

Bees were sampled on each plot four times between July 2019 to April 2020, one sampling within each of the four major climatic seasons of the area (i.e. November-December: short dry and cold season, March-April: long rainy and warm season, July: long dry and cold season, and September-October: short rainy and warm season). Bees were sampled by sweep netting and by using an Improved Prokopack aspirator (Model 1419, John W. Hock, Gainsville, Florida, USA). The sampling process was intensive, standardised and covered the active periods of bee foraging, that is from 09:00 to 17:00 EAT. During sampling, we moved slowly and gently in a parallel manner throughout each plot scouting for bees contacting the reproductive structures (anther and or stigma) of plants. All sampling was conducted during periods of no rains and strong winds. All sampled bees were directly frozen by transferring into a-18° motorable cooler (Waeco Coolfreeze CF-35, Dometic GmbH, Emsdetten, Germany) before onward transfer to a-80°C freezer in the laboratory at the International Centre of Insect Physiology and Ecology (*icipe*), Nairobi, Kenya. The direct freezing process helped in keeping the samples fresh, maintaining the structures of the bees for a longer time period, preventing bristles and damage while providing ease in handling during photo sessions. All bee specimens were identified to genus level following Michener (2007) and Eardley et al (2010) using a Zeiss^TM^ microscope affixed with an Axiocam 105 colour camera (Carl Zeiss microscope, Jena, Germany). Specimens were later sorted and assigned to species or morphospecies by Jane Machariafrom the National Museums of Kenya. We recorded all flowering plant species visited by bees on each study plot during each sampling round. During sweep netting walks, we emphasised on plant species in bloom as the main spots to locate bees. All plant species visited by bees were collected and identified to species level by an experienced plant taxonomist (Kennedy Matheka) at the National Museums of Kenya.

## Results

We recorded a total of 16124 bee visits on flowers, from 5 families, 43 genera and 181 species across the two counties. The most abundant bee family was the Apidae, making up 89.4% of all bee visits. This was followed by the families Halictidae (7.6%), Megachilidae (2.6%), Colletidae (0.3%) and Andrenidae (0.01%). At the county level, we recorded 4 out of the 5 families of bees known to exist in East Africa in Murang’a with the most visits from the family Apidae (91.7%) followed by Halictidae (6.8%), Megachilidae (1.3%) and Colletidae (0.2%). Meanwhile in Taita, we recorded 5 out of the 5 bee families with the most visits from the family Apidae (85%), followed by Halictidae (9.2%), Megachilidae (5.1%), Colletidae (0.63%) and Andrenidae (0.03%). Overall, the most abundant bee genus was *Apis*, making up 81.4% of all visits to flowers. This was followed by *Ceratina* (2.4%), *Lasioglossum* (2.2%), *Braunsapis* (1.8%), *Zonalictus* (1.5%), *Amegilla* (1.2%) and *Megachile* (1.1%). The least abundant genera were *Afromelecta* (0.01%), *Nomada* (0.01%), and *Thrinchostoma* (0.01%). In Murang’a, the genera *Apis* (87.4%), *Lasioglossum* (2.4%) and *Zonalictus* (1.6%) were the most abundant, while *Eupertersia* (0.01%) and *Nomada* (0.01%) were the least abundant. In Taita, the genera *Apis* (69.8%), *Ceratina* (4%), *Braunsapis* (3.2%) and *Amegilla* (3.1%) were the most abundant, while *Eupertersia* (0.02%) and *Thrinchostoma* (0.02%) were the least abundant. Across all counties, the most abundant species were *Apis mellifera* (Table S1).

We recorded bees visiting 312 plant species belonging to 175 genera and 55 families. In general, the most visited plant families were Asteraceae (24.4%), Lamiaceae (22.8%), Rubiaceae (19.2%), Capparaceae (7.3%), Acanthaceae (3.8%) and Fabaceae (3.2%). The least visited families were Ranunculaceae, Resedaceae, Scrophulariaceae and Polygalaceae accounting for 0.01% visitations each. At the county level, we recorded bees visiting 38 plant families in Murang’a with the most visited families being Rubiaceae (27.8%), Asteraceae (27.3%), Lamiaceae (21.6%) then Cyperaceae and Lauraceae with 3.6% each. The least visited plant families were Proteaceae, Ranunculaceae and Scrophulariaceae accounting for 0.01% of visits each. In Taita, we recorded bees visiting 40 plant families with the most visited families being Lamiaceae (25.1%), Capparaceae (21.1%), Asteraceae (18.9%), Fabaceae (5.4%) and Amaranthaceae (4.6%), while the least visited families were Brassicaceae, Ebenaceae, Meliaceae, Passifloraceae, Resedaceae and Polygalaceae, accounting for 0.02% of visits each.

Overall, the most visited plant genera were *Psychotria* (18%), *Ocimum* (10.6%), *Maerua* (7%), *Tithonia* (6.7%), *Lippia* (6.1%), *Bidens* (5.2%), *Justicia* (3.2%) and *Leucas* (2.4%). The least visited genera were *Veronica*, *Wahlenbergia*, *Zornia* and *Cyathula*, each accounting for 0.01% visitations respectively. At the county level, we recorded bees visiting 118 plant genera in Murang’a. The most visited plant genera were *Psychotria* (26.4%), *Tithonia* (10.1%), *Lippia* (9.1%), *Ocimum* (5.8%) and *Bidens* (5.5%), while the least visited genera were *Stylosanthes*, *Thunbergia*, *Veronica*, *Wahlenbergia* and *Zornia* with 0.01% of bee visitation on each. In Taita, we recorded bees visiting 104 plant genera and the most visited are *Maerua* (20.5%), *Ocimum* (19.9%), *Leucas* (5.9%), *Bidens* (4.5%) and *Conyza* (3.6%), while the least visited were *Spermacoce*, *Stylosanthes*, *Turraea*, *Urochloa* and *Cyathula*, each contributing 0.02% of bee visits. In general, the most visited plant species were *Psychotria mahonii* (17.4%), *Ocimum gratissimum* (7.5%), *Tithonia diversifolia* (6.7%), *Maerua sp.1* (6.5%), *Lippia javanica* (6%), *Bidens Pilosa* (4.6%), *Persea sp.1* (2.4%), *Kyllinga brevifolia* (2.3%), *Leucas grandis* (2.2%), and *Justicia calyculata* (2.1%). The least visited plant species were, *Thunbergia alata*, *Turrea robusta*, *Vernonia auriculifera*, *Veronica abyssinica*, *Wahlenbergia abyssinica* and *Zornia setosa*, each accounting for 0.01% visitation. At the county level, we recorded bee visits on 179 plant species in Murang’a. The most visited plant species were *Psychotria mahonii* (26.5%), *Tithonia diversifolia* (10.1%), *Lippia javanica* (9.1%), *Bidens spilosa* (5.5%) and *Ocimum gratissimum* (4.3%). The least visited plant species were *Stylosanthes fruticosa*, *Thunbergia alata*, *Veronica abyssinica*, *Wahlenbergia abyssinica* and *Zornia setosa*, each accounting for 0.01% visitation. In Taita, we recorded bees on 175 plant species. The most visited plant species were *Maerua sp. 1* (18.9%), *Ocimum gratissimum* (13.5%), *Leucas grandis* (5.4%), *Conyza newii* (5.3%), *Bidens pilosa* and *Digera muricata* with 2.7% of bee visits each. The least visited species were *Spermacoce pusilla*, *Stylosanthes fruticosa*, *Turraea robusta*, *Urochloa trichopus*, *Vernonia auriculifera* and *Cyathula orthacantha*, each accounting for 0.02% of total plants visited (Table S2).

## Discussion

We here present evidence on species assemblage patterns and pictorial data of bee pollinators and the plants they visit in two counties (Murang’a and Taita-Taveta) of Kenya. We believe these photos will improve on the knowledge and awareness of the different plants and bee visitors in the local communities of these two counties and might further serve as an important resource for future studies that would require community participation with local indigenes. Our results showed marked differences in bee community assemblage patterns between the two counties. Bee abundance was significantly more in Murang’a than in Taita. This might be due to the huge number of honeybees deployed in hundreds of managed bee hives spread across the highlands of the Aberdare forests (Dzekashu et al., 2023). On the other hand, bee species richness was more in the warm lowlands of Taita than in Murang’a. This suggests that the warm conditions of the lower elevations of Taita provides the ideal thermoregulatory requirements and high seasonal turnover of floral resources for bees (Dzekashu et al., 2022). Nonetheless, the number of visited plant species did not differ across the two counties. Though just 13% of the bee-visited plant species overlapped across both counties, they had an almost equal number of unique species between them (137 species in Muanga and 134 species in Taita). This indicates that variations in ecological factors (e.g. climate seasonality) within these counties might be playing important roles in supporting the observed distinct assemblage patterns of plant species (Dzekashu et al., 2022).

Studies aimed at examining global change influences on species assemblage patterns with community participation (through citizen science) has led to the emergence of large-scale data sets with relative high inferential power (Rondeau et al., 2023). Such studies typically requires hypotheses formulations, data collection and assessment of ecological trends (Brown & Williams, 2019). Therefore citizen science has the ability to enhance the scope of ecological data collection, contribute immensely to scientific awareness, social interactions and with an added advantage for participants when they relate with nature (Mason & Arathi, 2019). While citizen science can produce large data sets with high potentials for errors (especially in species identification) and biases in experimental set up and data collection in ecology (Dickinson et al., 2010), such errors can be minimised through the implementation of rigorious experemental designs (and expert’s identified species photos), while experimental bias can be mitigated through awareness trainings on the importance of maintaining proper scientific standards and upholding data integrity at all times. These errors and biases can also be controlled in circumstances whereby the collected data is entered into a centralised database for inspection and vetoing by experts before publishing. Moreover, in ecological studies, the study design and not volunteer involvement could be a limiting factor on the integrity of data collected (Brown & Williams, 2019). This could be actual for observational studies producing stronger inferences when designed with a well-structured a priori hypotheses and considerable investments in efforts and time in training and regularly monitoring the performance of volunteers (Brown & Williams, 2019; Nichols et al., 2012; Dickinson et al., 2010).

Recent technological advancements including the development of portable devices such as digital cameras and smartphones with applications capable of handling ecological informations has revolutionalised the process of environmental data gathering with perfect detectibility in species observation studies (Sheard et al., 2024). These portable devices have enhanced the collection, digitalization and uploading or submission of species occurences data into online repositories such as the Global Biodiversity Information Facility (GBIF), iNaturalist, Pl@ntNet, Encyclopedia Of Life (EOL), iSpot, etc. Though not exhaustive, our photos presented in this manuscript informs on the most common plant and bee visitors and could serve as important guides or starting ground for studies involving the digitalisation of plant-bee pollinator interactions in the Eastern Afromontane Biodiversity Hotspots of Kenya (EABH).

In this study, we explore the usefullness of images in ecological studies. Using digitalised photograghs, we elaborated on the community assemblage patterns of plants and their bee pollinators in the EABH of Kenya and further discussed on the importance of exploiting images for research in biodiversity and conservation. We opined that photographs can ease understanding of species community assemblage patterns and can also encourage individuals (through citizen science projects), communities and policy makers to enact decisions that would advance conservation efforts. A huge challenge in biodiversity is the time, effort and cost associated with the collection and processing of samples (particularly taxonomic identifications). The ever-increasing willingness of individual volunteers (as citizen scientist) to participate in biodiversity studies can help increase sampling efforts when properly trained on standard scientific protocols. The presence of digital images can go a long way to inspire participation in sample sorting and processing (with the vetting of taxonomists), leading to a gain in time while reducing cost. In the tropics, several new species are being discovered, but there is limited and at times non-existent archived data sources, making it difficult to compare current changes in species communities to the past. Therefore, technological advances that helps in digitalising species through photography can be great tools for future biodiversity studies.

## Supporting information

https://figshare.com/s/ff5f1c01bfc24c6db94e

## Acknowledgements

We gratefully acknowledge the financial support for this research by the following organizations and agencies: JRS Biodiversity Foundation (grant number: 60610), UK’s Foreign, Commonwealth & Development Office (FCDO), the Swedish International Development Cooperation Agency (Sida), the Swiss Agency for Development and Cooperation (SDC), the Federal Democratic Republic of Ethiopia, the Government of the Republic of Kenya, and the South African National Research Foundation Incentive funding for Rated researchers to AAY and CWWP. Fairo Dzekashu was supported by a German Academic Exchange Services (DAAD) funded Ph. D. scholarship in the framework of the African Regional Postgraduate Programme in Insect Science (ARPPIS). We thank Alain Pauly for his comments on the identified bee species. We are grateful to Kimani James Kinuthia, Ephantus Kimani, James N. Kimani, Kika Junior, Benson Mwakachola, Darius Mwambala, and Mwadime Mjomba for their assistance in the various stages of field work.

## References

Alan Irwin. (1995). Citizen science: A study of people, excercise and sustainable development. In Environment and Society. Routledge.

Baldock, K. C. R., Memmott, J., Carlos Ruiz-Guajardo, J., Roze, D., & Stone, G. N. (2011). Daily temporal structure in African savanna flower visitation networks and consequences for network sampling. Ecology, 92(3), 687–698. 10.1890/10-1110.1

Bates, D., Mächler, M., Bolker, B., & Walker, S. (2015). Fitting Linear Mixed-Effects Models Using lme4. Journal of Statistical Software, 67(1). 10.18637/jss.v067.i01

Bonney, R., Cooper, C. B., Dickinson, J., Kelling, S., Phillips, T., Rosenberg, K. V., & Shirk, J. (2009). Citizen science: A developing tool for expanding science knowledge and scientific literacy. BioScience, 59(11), 977–984. 10.1525/bio.2009.59.11.9

Bonney, R., Shirk, J. L., Phillips, T. B., Wiggins, A., Ballard, H. L., Miller-Rushing, A. J., & Parrish, J. K. (2014). Next steps for citizen science. Science, 343(6178), 1436–1437. 10.1126/science.1251554

Brown, E. D., & Williams, B. K. (2019). The potential for citizen science to produce reliable and useful information in ecology. Conservation Biology, 33(3), 561–569. 10.1111/cobi.13223

Depauw, L., Blondeel, H., De Lombaerde, E., De Pauw, K., Landuyt, D., Lorer, E., Vangansbeke, P., Vanneste, T., Verheyen, K., & De Frenne, P. (2022). The use of photos to investigate ecological change. Journal of Ecology, 110(6), 1220–1236. 10.1111/1365-2745.13876

Dickinson, J. L., Zuckerberg, B., & Bonter, D. N. (2010). Citizen science as an ecological research tool: Challenges and benefits. *Annual Review of Ecology*, Evolution, and Systematics, 41, 149–172. 10.1146/annurev-ecolsys-102209-144636

Dormann, C. F., Fründ, J., & Schaefer, H. M. (2017). Identifying Causes of Patterns in Ecological Networks: Opportunities and Limitations. Annual Review of Ecology, Evolution, and Systematics, 48, 559–584. 10.1146/annurev-ecolsys-110316-022928

Dzekashu, F. F., Peters, M. K., SteffanLDewenter, I., Macharia, J. M., Matheka, K. W., Yusuf, A. A., Pirk, C. W. W., & Lattorff, H. M. G. (2025). Plants and Bee Visitors in the Eastern Afromontane Biodiversity Hotspot of Kenya, East Africa. The Bulletin of the Ecological Society of America, 106(1). 10.1002/bes2.2211

Dzekashu, F. F., Pirk, C. W. W., Yusuf, A. A., Classen, A., Kiatoko, N., SteffanLDewenter, I., Peters, M. K., & Lattorff, H. M. G. (2023). Seasonal and elevational changes of plantLpollinator interaction networks in East African mountains. Ecology and Evolution, 13(5), 1–16. 10.1002/ece3.10060

Dzekashu, F. F., Yusuf, A. A., Pirk, C. W. W., Steffan-Dewenter, I., Lattorff, H. M. G., & Peters, M. K. (2022). Floral turnover and climate drive seasonal bee diversity along a tropical elevation gradient. Ecosphere, 13(3). 10.1002/ecs2.3964

Dzekashu, F. F., Yusuf, A. A., Takemoto, K., Peters, M. K., Lattorff, H. M. G., Steffan-Dewenter, I., & Pirk, C. W. W. (2024). Network resilience of plant-bee interactions in the Eastern Afromontane Biodiversity Hotspot. Ecological Indicators, 166, 112415. 10.1016/j.ecolind.2024.112415

Eardley, C., Kuhlmann, M., & Pauly, A. (2010). The bee genera and subgenera of sub-Saharan Africa. Belgian Development Cooperation Brussels.

Eilers, E. J., Kremen, C., Smith Greenleaf, S., Garber, A. K., & Klein, A.-M. (2011). Contribution of Pollinator-Mediated Crops to Nutrients in the Human Food Supply. PLoS ONE, 6(6), e21363. 10.1371/journal.pone.0021363

ElQadi, M. M., Dorin, A., Dyer, A., Burd, M., Bukovac, Z., & Shrestha, M. (2017). Mapping species distributions with social media geo-tagged images: Case studies of bees and flowering plants in Australia. Ecological Informatics, 39, 23–31. 10.1016/j.ecoinf.2017.02.006

Gebrechorkos, S. H., Hülsmann, S., & Bernhofer, C. (2019). Changes in temperature and precipitation extremes in Ethiopia, Kenya, and Tanzania. International Journal of Climatology, 39(1), 18–30. 10.1002/joc.5777

Gonella, P. M., Rivadavia, F., & Fleischmann, A. (2015). Drosera magnifica (Droseraceae): The largest new world sundew, discovered on Facebook. Phytotaxa, 220(3), 257–267. 10.11646/phytotaxa.220.3.4

Guy, T. J., Hutchinson, M. C., Baldock, K. C. R., Kayser, E., Baiser, B., Staniczenko, P. P. A., Goheen, J. R., Pringle, R. M., & Palmer, T. M. (2021). Large herbivores transform plant-pollinator networks in an African savanna. Current Biology, 31(13), 2964–2971.e5. 10.1016/j.cub.2021.04.051

Hartig, F. (2016). DHARMa: Residual Diagnostics for Hierarchical (Multi-Level / Mixed) Regression Models. In CRAN: Contributed Packages. 10.32614/CRAN.package.DHARMa

Lanuza, J. B., Knight, T. M., Montes-Perez, N., Glenny, W., Acuña, P., Albrecht, M., Artamendi, M., Badenhausser, I., Bennett, J. M., Biella, P., Bommarco, R., Cappellari, A., Castro, S., Clough, Y., Colom, P., Costa, J., Cyrille, N., de Manincor, N., Dominguez-Lapido, P.,…Bartomeus, I. (2025). EuPPollNet: A European Database of Plant-Pollinator Networks. Global Ecology and Biogeography, 34(2). 10.1111/geb.70000

Lasway, J. V., Steffan-Dewenter, I., Njovu, H. K., Kinabo, N. R., Eardley, C., Pauly, A., & Peters, M. K. (2022). Positive effects of low grazing intensity on East African bee assemblages mediated by increases in floral resources. Biological Conservation, 267, 109490. 10.1016/j.biocon.2022.109490

Legendre, P., & Legendre, L. (2012). Complex ecological data sets. Developments in Environmental Modelling. 10.1016/B978-0-444-53868-0.50001-0

Mason, L., & Arathi, H. S. (2019). Assessing the efficacy of citizen scientists monitoring native bees in urban areas. Global Ecology and Conservation, 17, e00561. 10.1016/j.gecco.2019.e00561

Mertens, J. E. J., Brisson, L., Janeček, Š., Klomberg, Y., Maicher, V., Sáfián, S., Delabye, S., Potocký, P., Kobe, I. N., Pyrcz, T., & Tropek, R. (2021). Elevational and seasonal patterns of butterflies and hawkmoths in plant-pollinator networks in tropical rainforests of Mount Cameroon. Scientific Reports, 11(1), 1–12. 10.1038/s41598-021-89012-x

Michener, C. D. (2007). The bees of the world (Issue 595.799 M53/2007).

Nichols, J. D., Cooch, E. G., Nichols, J. M., & Sauer, J. R. (2012). Studying biodiversity: Is a new paradigm really needed? BioScience, 62(5), 497–502. 10.1525/bio.2012.62.5.11

Oksanen, A. J., Blanchet, F. G., Kindt, R., Legen-, P., Minchin, P. R., Hara, R. B. O., Simpson, G. L., Solymos, P.,&Stevens, M. H. H. (2018). Community Ecology Package. …Ecology Package…, 263. http://mirror.bjtu.edu.cn/cran/web/packages/vegan/vegan.pdf

Ollerton, J., Winfree, R., & Tarrant, S. (2011). How many flowering plants are pollinated by animals? Oikos, 120(3), 321–326. 10.1111/j.1600-0706.2010.18644.x

Orodho, A. B. (2006). Country pasture/forage resource profiles. FAO/Republic of Kenya.

Potts, S. G., Imperatriz-Fonseca, V., Ngo, H. T., Aizen, M. A., Biesmeijer, J. C., Breeze, T. D., Dicks, L. V., Garibaldi, L. A., Hill, R., Settele, J., & Vanbergen, A. J. (2016). Safeguarding pollinators and their values to human well-being. Nature, 540(7632), 220–229. 10.1038/nature20588

Powney, G. D., Carvell, C., Edwards, M., Morris, R. K. A., Roy, H. E., Woodcock, B. A., & Isaac, N. J. B. (2019). Widespread losses of pollinating insects in Britain. Nature Communications, 10(1), 1–6. 10.1038/s41467-019-08974-9

R Core Team. (2024). R: A Language and Environment for Statistical Computing_. R Foundation for Statistical Computing, Vienna, Austria. <https://www.R-project.org/>. https://www.r-project.org/

Rondeau, S., Gervais, A., Leboeuf, A., Drapeau Picard, A. P., Larrivée, M., & Fournier, V. (2023). Combining community science and taxonomist expertise for large-scale monitoring of insect pollinators: Perspective and insights from Abeilles citoyennes. Conservation Science and Practice, 5(10), 1–15. 10.1111/csp2.13015

Sanseverino, M. E., Whitney, M. J., & Higgs, E. S. (2016). Exploring Landscape Change in Mountain Environments with the Mountain Legacy Online Image Analysis Toolkit. Mountain Research and Development, 36(4), 407–416. 10.1659/MRD-JOURNAL-D-16-00038.1

Sheard, J. K., Adriaens, T., Bowler, D. E., Büermann, A., Callaghan, C. T., Camprasse, E. C. M., Chowdhury, S., Engel, T., Finch, E. A., von Gönner, J., Hsing, P.-Y., Mikula, P., Rachel Oh, R. Y., Peters, B., Phartyal, S. S., Pocock, M. J. O., Wäldchen, J., & Bonn, A. (2024). Emerging technologies in citizen science and potential for insect monitoring. Philosophical Transactions of the Royal Society B: Biological Sciences, 379(1904). 10.1098/rstb.2023.0106

Stein, K., Coulibaly, D., Balima, L. H., Goetze, D., Linsenmair, K. E., Porembski, S., Stenchly, K., & Theodorou, P. (2021). Plant-pollinator networks in savannas of Burkina Faso, West Africa. Diversity, 13(1), 1–14. 10.3390/d13010001

Vanbergen, A. J., Garratt, M. P., Vanbergen, A. J., Baude, M., Biesmeijer, J. C., Britton, N. F., Brown, M. J. F., Brown, M., Bryden, J., Budge, G. E., Bull, J. C., Carvell, C., Challinor, A. J., Connolly, C. N., Evans, D. J., Feil, E. J., Garratt, M. P., Greco, M. K., Heard, M. S.,…Wright, G. A. (2013). Threats to an ecosystem service: Pressures on pollinators. Frontiers in Ecology and the Environment, 11(5), 251–259. 10.1890/120126

Woodard, S. H., Federman, S., James, R. R., Danforth, B. N., Griswold, T. L., Inouye, D., McFrederick, Q. S., Morandin, L., Paul, D. L., Sellers, E., Strange, J. P., Vaughan, M., Williams, N. M., Branstetter, M. G., Burns, C. T., Cane, J., Cariveau, A. B., Cariveau, D. P., Childers, A.,…Wehling, W. (2020). Towards a U.S. national program for monitoring native bees. Biological Conservation, 252(November). 10.1016/j.biocon.2020.108821

