## Supplementary material for "A citizen science photo guide of plants and bee visitors in the Eastern Afromontane Biodiversity Hotspot of Kenya": https://figshare.com/s/ff5f1c01bfc24c6db94e

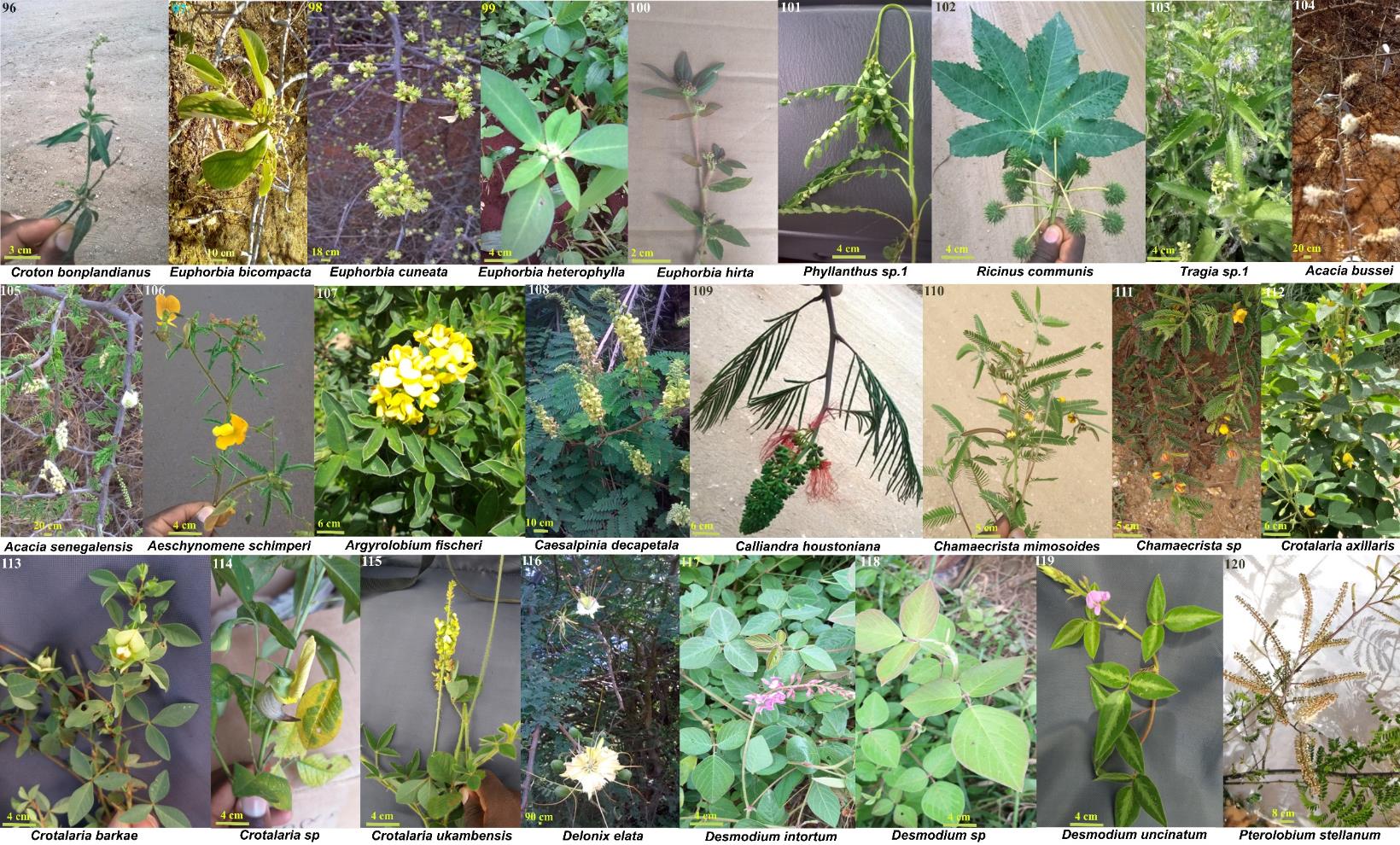

**Figure S1**. Plant species on which bee species were found visiting on study plots. Plant species are arranged according to families; 96-103: Euphorbiaceae, 104-120: Fabaceae. Measurement in cm shows the scale of each photo. Photo credit: Fairo Dzekashu

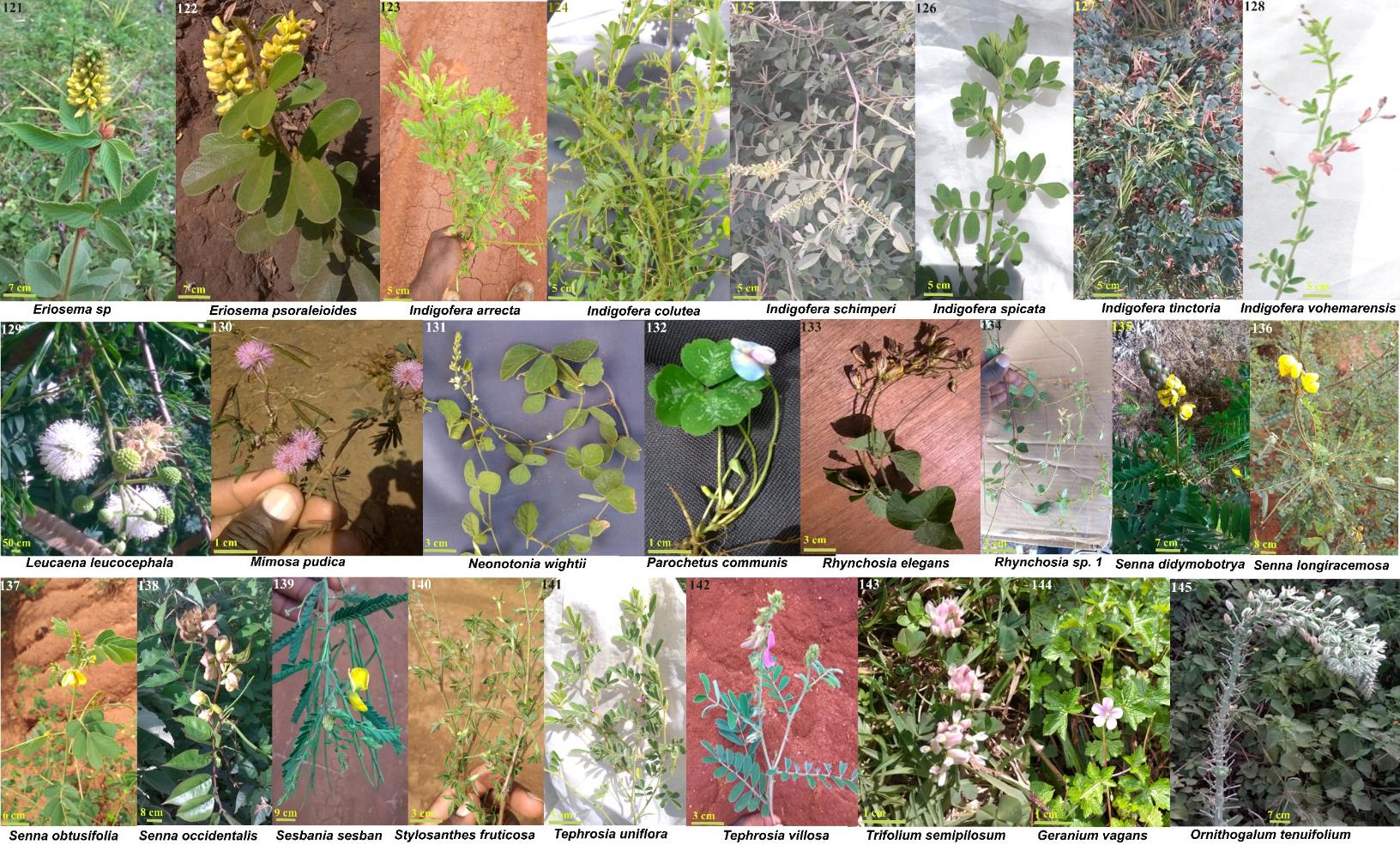

**Figure S2**. Plant species on which bee species were found visiting on study plots. Plant species are arranged according to families; 121-143: Fabaceae, 144: Geraniaceae, 145: Hyacinthaceae. Measurement in cm shows the scale of each photo. Photo credit: Fairo Dzekashu

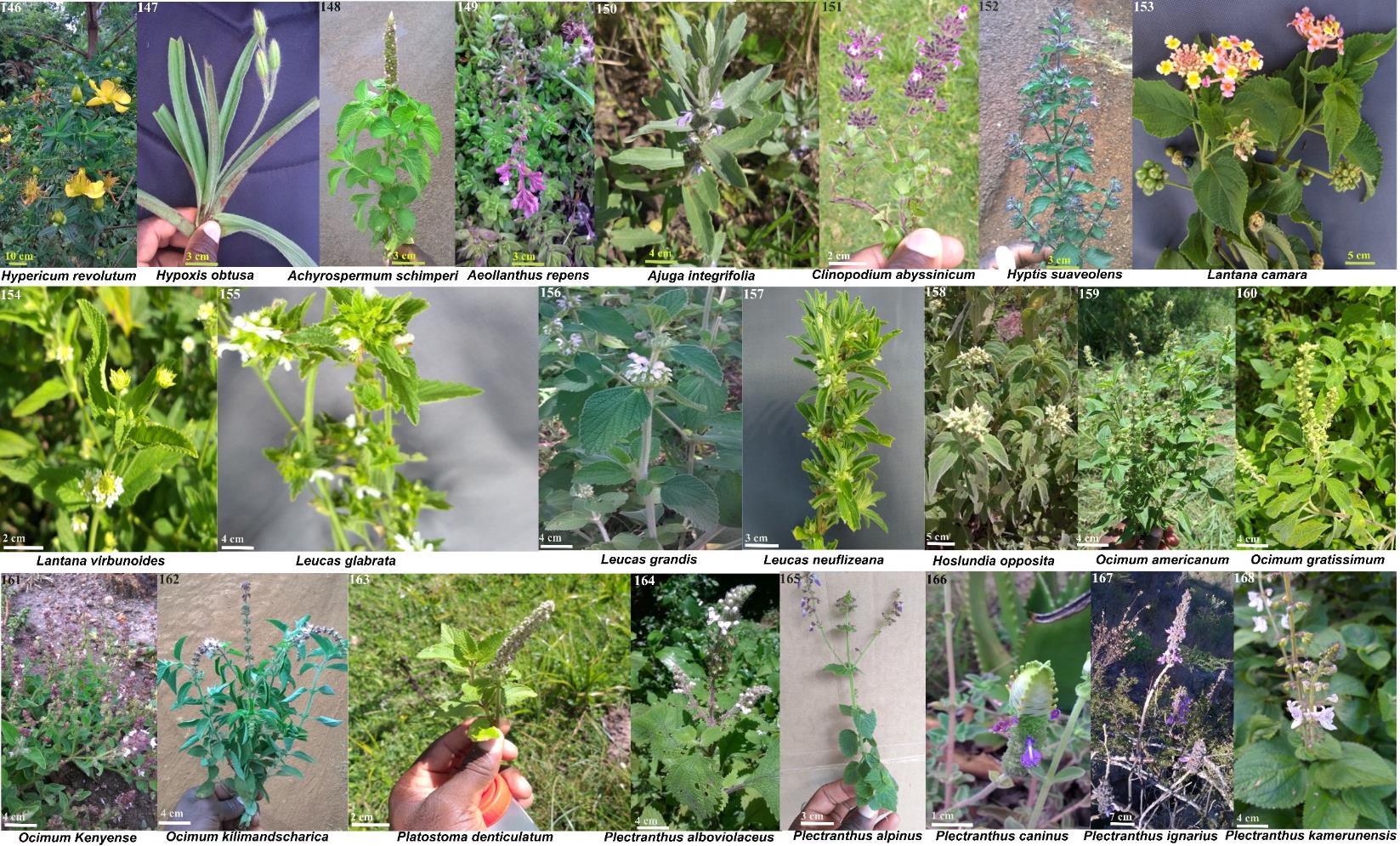

**Figure S3**. Plant species on which bee species were found visiting on study plots. Plant species are arranged according to families; 146: Hypericaceae, 147: Hypoxidaceae, 148-168: Lamiaceae. Measurement in cm represents the scale of each photo. Photo credit: Fairo Dzekashu

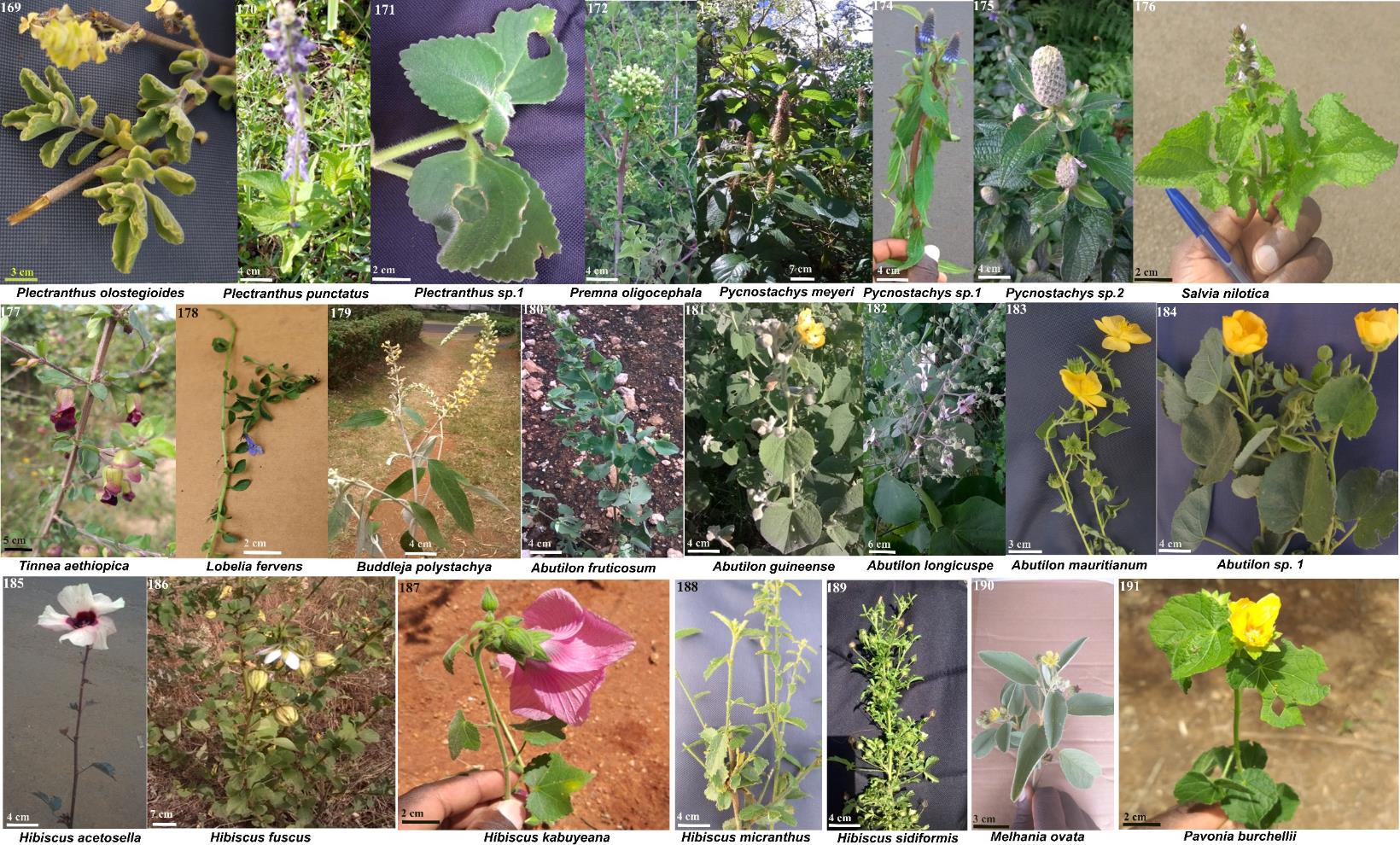

**Figure S4**. Plant species on which bee species were found visiting on study plots. Plant species are arranged according to families; 169-177: Lamiaceae, 178: Lobeliaceae, 179: Loganiaceae, 180-191: Malvaceae. Measurement in cm indicates the scale of each photo. Photo credit: Fairo Dzekashu

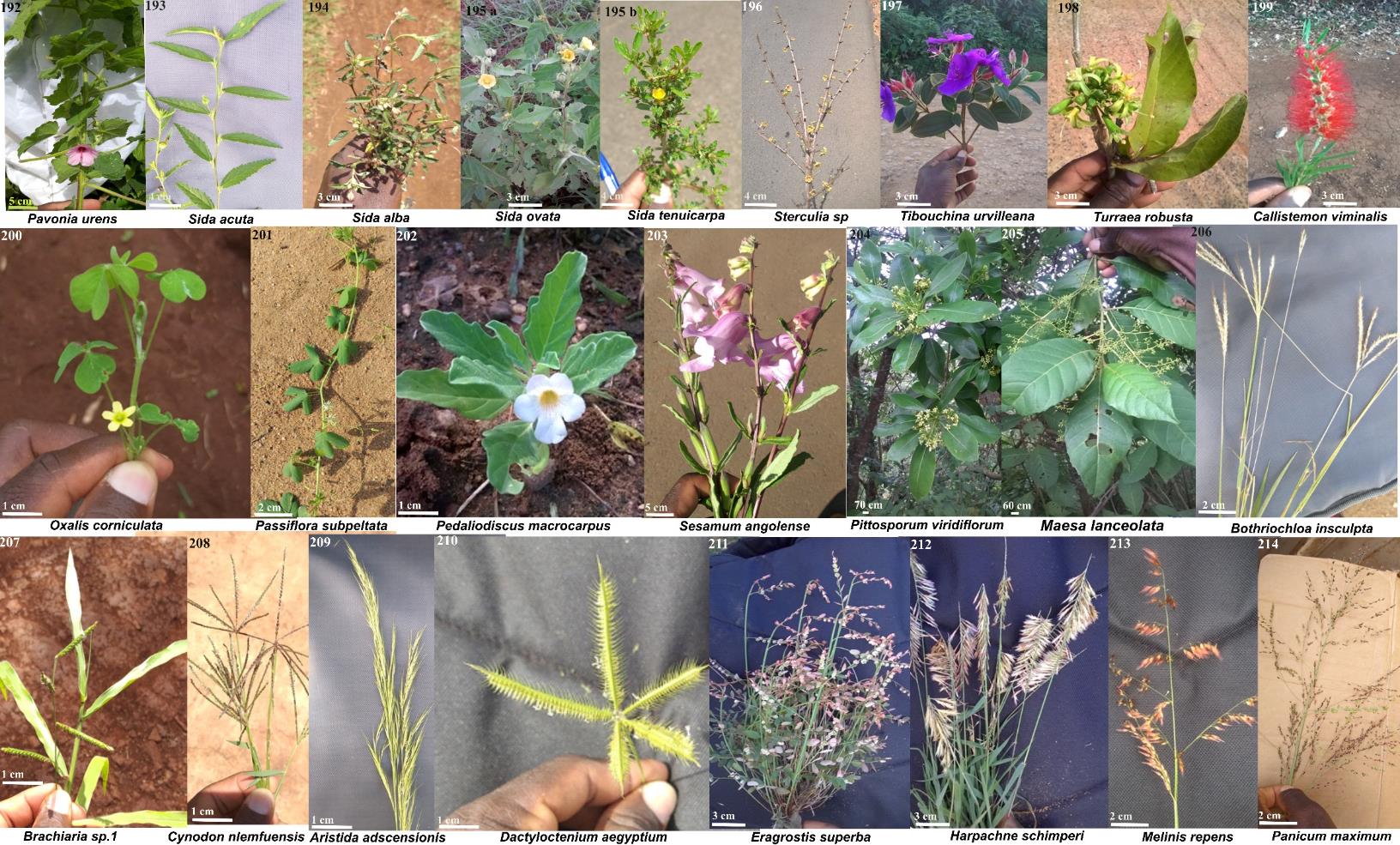

**Figure S5**. Plant species on which bee species were found visiting on study plots. Plant species are arranged according to families; 192-196: Malvaceae, 197: Melastomataceae, 198: Meliaceae, 199: Myrtaceae, 200: Oxalidaceae, 201: Passifloraceae, 202-203: Pedaliaceae, 204: Pittosporaceae, 205: Primulaceae, 206-214: Poaceae. Measurement in cm shows the scale of each photo. Photo credit: Fairo Dzekashu

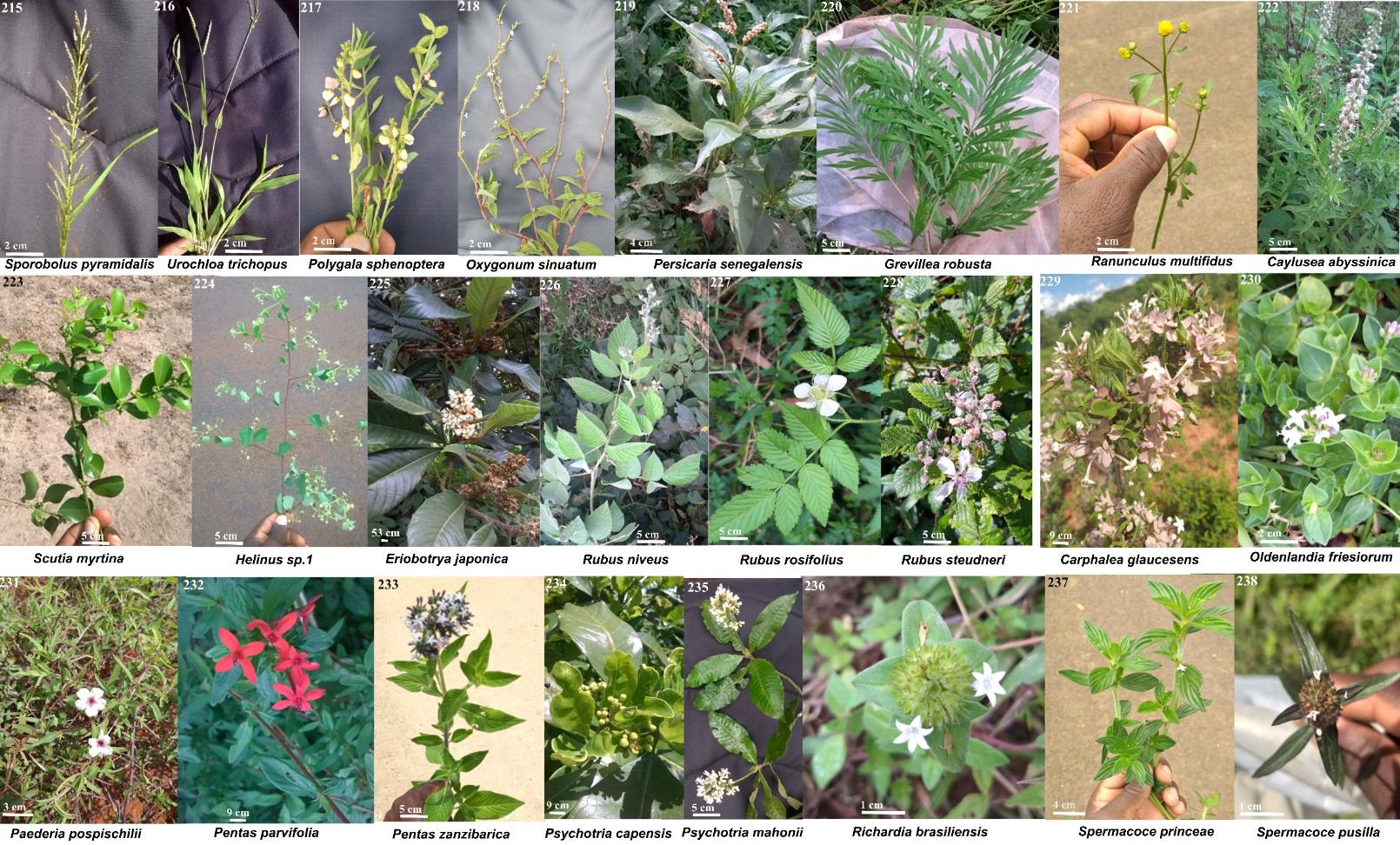

**Figure S6**. Plant species on which bee species were found visiting on study plots. Plant species are arranged according to families; 215-216: Poaceae, 217-219: polygalaceae, 220: Proteaceae, 221: Ranunculaceae, 222: Resedaceae, 223-224: Rhamnaceae, 225-238: Rubiaceae. Measurement in cm represents the scale of each photo. Photo credit: Fairo Dzekashu

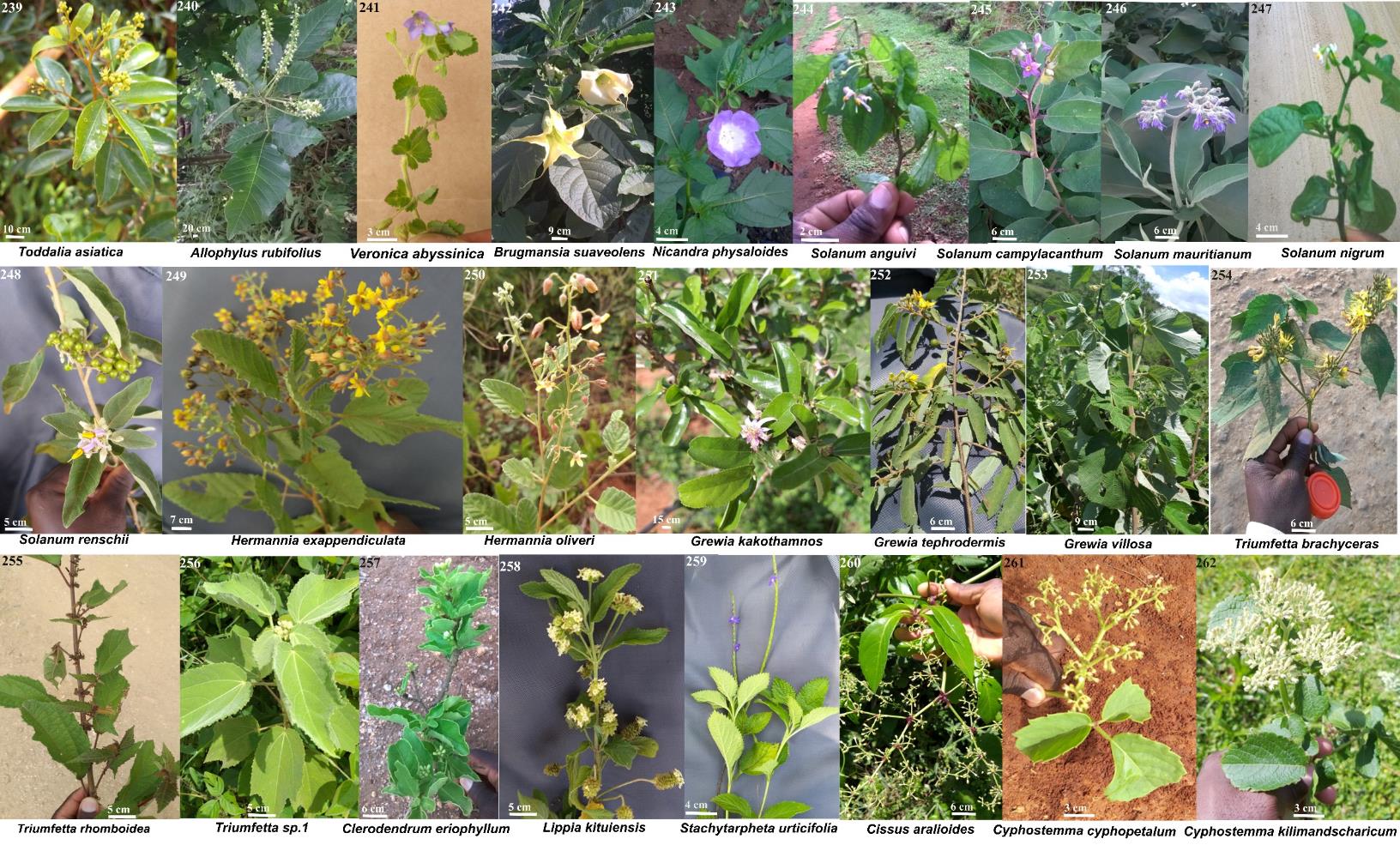

**Figure S7**. Plant species on which bee species were found visiting on study plots. Plant species are arranged according to families; 239: Rutaceae, 240: Sapindaceae, 241: Scrophulariaceae, 242-248: Solanaceae, 249-250: Sterculiaceae, 251-257: Tiliaceae, 257-259: Verbenaceae, 260-262: Vitaceae. Photo credit: Fairo Dzekashu

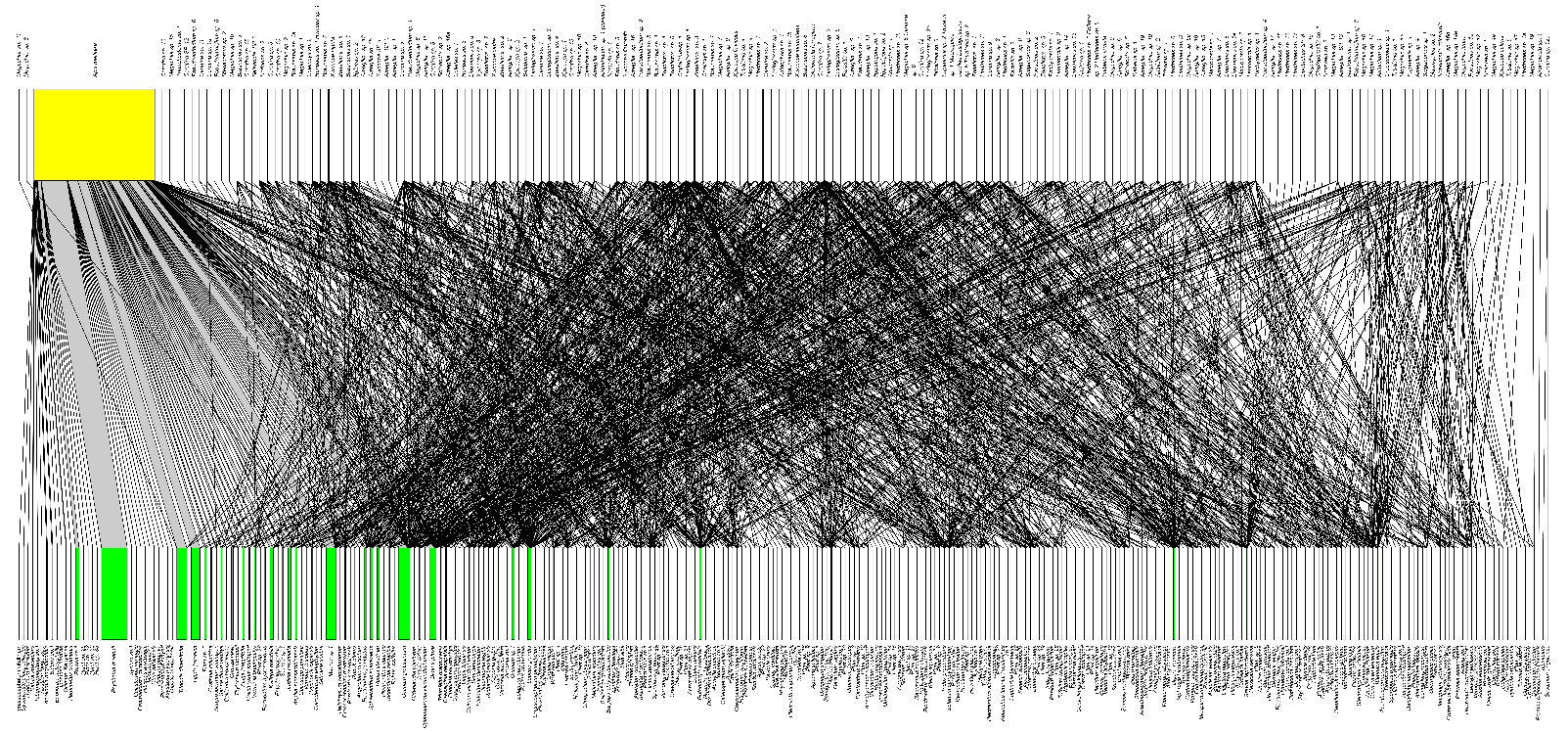

**Figure S8.** Bipartite graph of Plant-bee interactions of all plants and bee species recorded in the two counties (Muranga and Taita). Bee pollinator species (yellow vertical bars at the top) recorded visiting plant species (green vertical bars at the bottom). Grey lines indicate links between bee and plant species. The width of each vertical bar indicates visitation frequency of each species, while the size of the lines indicates interaction strength of each species.

**Table S1:** Differences in community assemblage patterns (Bee Abundance (with HB), Bee Abundance (no HB), Bee_Species, Plant_Species) between the two counties (Taita and Muranga. Analysed using generalized mixed models (GLMMs) with a negative binomial. Model outputs: standard error (SE), standard deviation (z value), approximate significance of terms (P-value). Asterisks denote significant codes (*: <0.05, **: <0.01, ***: <0.001).

| **Response** | **Predictor** | **Estimate** | **SE** | **z value** | **P-value** |  |
| --- | --- | --- | --- | --- | --- | --- |
| Bee Abundance (with HB) | (Intercept) | 5.5662 | 0.2233 | 24.926 | < 2e-16 | *** |
|  | county | -0.8331 | 0.3173 | -2.625 | **0.00865** | ****** |
| Bee Abundance (no HB) | (Intercept) | 3.7154 | 0.1602 | 23.187 | <2e-16 | *** |
|  | county | 0.3041 | 0.2254 | 1.349 | 0.177 |  |
| Bee_Species | (Intercept) | 2.9219 | 0.1064 | 27.46 | <2e-16 | *** |
|  | county | 0.3401 | 0.1473 | 2.31 | **0.0209** | ***** |
| Plant_Species | (Intercept) | 2.70053 | 0.0886 | 30.481 | <2e-16 | *** |
|  | county | -0.09337 | 0.12531 | -0.745 | 0.456 |  |

**Table 2:** Bee species list, the elevational distribution of bees and the percentage abundance per species across the two counties. Non-metric multidimensional scaling ordination distances (NMDS1 & NMDS 2), coefficient of variation (**R^2^**), significance in ordination spaces by intrinsic species (P-value & Sig.), Min_Ele (Minimum elevation of species occurrence), Max_Ele (Maximum elevation of species occurrence), yellow colours (Species occurrence in Taita only), green colours (Species occurrence in Muranga only), blue colours (Species occurrence in Taita and Muranga), % abun (percentage abundance of each species).

| **Bee species** | **NMDS1** | **NMDS2** | **R^2^** | **P-value** | **Sig.** | **Min_Ele** | **Max_Ele** | **% abun** |
| --- | --- | --- | --- | --- | --- | --- | --- | --- |
| *Acunomia sp. 1* | -0.34406 | -0.93895 | 0.0217 | 0.623 |  | 679 | 1462 | 0.030 |
| *Afranthidium sp. 1 (concolor)* | 0.73248 | -0.68078 | 0.0183 | 0.557 |  | 1632 | 1632 | 0.006 |
| *Afromelecta sp. 1* | -0.52065 | -0.85377 | 0.0711 | 0.226 |  | 730 | 730 | 0.006 |
| *Amegilla sp. 1* | 0.03694 | 0.99932 | 0.0188 | 0.507 |  | 1544 | 1633 | 0.006 |
| *Amegilla sp. 2* | -0.06524 | -0.99787 | 0.085 | 0.104 |  | 627 | 1864 | 0.054 |
| ***Amegilla sp. 3*** | **-0.25327** | **-0.9674** | **0.2374** | **0.006** | ****** | **529** | **1695** | 0.431 |
| ***Amegilla sp. 4*** | **-0.73444** | **-0.67867** | **0.1649** | **0.021** | ***** | **627** | **1831** | 0.072 |
| ***Amegilla sp. 5*** | **-0.3639** | **-0.93144** | **0.1883** | **0.013** | ***** | **627** | **1539** | 0.066 |
| *Amegilla sp. 6* | -0.80535 | 0.5928 | 0.0153 | 0.692 |  | 730 | 1695 | 0.090 |
| *Amegilla sp. 6b* | -0.35388 | -0.93529 | 0.0836 | 0.166 |  | 1053 | 1053 | 0.006 |
| *Amegilla sp. 7* | -0.32911 | -0.94429 | 0.0947 | 0.078 | . | 730 | 1671 | 0.114 |
| *Amegilla sp. 8* | -0.91572 | -0.40181 | 0.0759 | 0.206 |  | 529 | 529 | 0.006 |
| *Amegilla sp. 9* | 0.09806 | -0.99518 | 0.065 | 0.211 |  | 981 | 1632 | 0.078 |
| ***Amegilla sp. 10*** | **-0.62299** | **-0.78223** | **0.1411** | **0.03** | ***** | **529** | **1053** | 0.012 |
| ***Amegilla sp. 11*** | **-0.28296** | **-0.95913** | **0.2047** | **0.019** | ***** | **981** | **1053** | 0.036 |
| *Amegilla sp. 12* | -0.22629 | -0.97406 | 0.1206 | 0.11 |  | 981 | 981 | 0.006 |
| *Amegilla sp. 13* | 0.29023 | -0.95696 | 0.0337 | 0.445 |  | 895 | 1864 | 0.012 |
| *Amegilla sp. 14* | -0.31129 | -0.95031 | 0.0833 | 0.121 |  | 740 | 1630 | 0.030 |
| *Amegilla sp. 15* | 0.18077 | 0.98352 | 0.0042 | 0.931 |  | 1218 | 1695 | 0.018 |
| *Amegilla sp. 16* | -0.1009 | -0.9949 | 0.0821 | 0.136 |  | 1053 | 1864 | 0.018 |
| ***Amegilla sp. 16a*** | **-0.28296** | **-0.95913** | **0.2047** | **0.019** | ***** | **981** | **1053** | 0.012 |
| *Amegilla sp. 16b* | -0.22629 | -0.97406 | 0.1206 | 0.11 |  | 981 | 981 | 0.006 |
| *Amegilla sp. 17* | -0.5377 | -0.84314 | 0.1263 | 0.093 | . | 679 | 679 | 0.006 |
| *Amegilla sp. 18* | 0.33284 | -0.94298 | 0.0462 | 0.336 |  | 981 | 1700 | 0.066 |
| *Amegilla sp. 19* | -0.3519 | -0.93604 | 0.1665 | 0.053 | . | 526 | 526 | 0.006 |
| *Andrena sp. 1* | 0.38654 | 0.92227 | 0 | 0.999 |  | 679 | 1695 | 0.012 |
| *Anthidium sp. 1* | -0.91572 | -0.40181 | 0.0759 | 0.206 |  | 529 | 529 | 0.012 |
| *Anthidium sp. 2* | -0.19284 | 0.98123 | 0.0544 | 0.295 |  | 1528 | 1528 | 0.006 |
| *Anthophora sp. 1* | -0.53281 | -0.84623 | 0.0507 | 0.272 |  | 679 | 1352 | 0.012 |
| ***Apis mellifera*** | **0.80579** | **-0.5922** | **0.5469** | **0.001** | ******* | **529** | **2528** | 81.434 |
| *Austronomia sp. 1* | -0.6809 | -0.73237 | 0.0022 | 0.962 |  | 671 | 1831 | 0.018 |
| *Braunsapis sp. 1* | 0.8835 | -0.46844 | 0.087 | 0.125 |  | 981 | 2192 | 0.221 |
| *Braunsapis sp. 1a* | -0.0086 | -0.99996 | 0.1066 | 0.063 | . | 627 | 1563 | 0.036 |
| *Braunsapis sp. 1b* | 0.36407 | -0.93137 | 0.0182 | 0.7 |  | 659 | 1352 | 0.024 |
| *Braunsapis sp. 2* | 0.12491 | -0.99217 | 0.0657 | 0.207 |  | 659 | 2528 | 0.574 |
| *Braunsapis sp. 3* | 0.66838 | 0.74382 | 0.0218 | 0.607 |  | 529 | 2390 | 0.138 |
| *Braunsapis sp. 4* | 0.07183 | -0.99742 | 0.0769 | 0.163 |  | 679 | 2192 | 0.108 |
| *Braunsapis sp. 5* | -0.26172 | -0.96514 | 0.0741 | 0.155 |  | 526 | 2070 | 0.688 |
| ***Braunsapis sp. 6*** | **-0.48683** | **-0.8735** | **0.2109** | **0.005** | ****** | **627** | **1544** | 0.054 |
| *Braunsapis sp. 7* | 0.36644 | -0.93044 | 0.007 | 0.805 |  | 669 | 2070 | 0.036 |
| *Ceratina sp. 1* | 0.75739 | -0.65297 | 0.0332 | 0.455 |  | 659 | 2135 | 0.144 |
| *Ceratina sp. 1a* | 0.97407 | -0.22626 | 0.0158 | 0.727 |  | 1218 | 2135 | 0.018 |
| *Ceratina sp. 2* | -0.30427 | -0.95259 | 0.0565 | 0.241 |  | 659 | 1700 | 0.520 |
| *Ceratina sp. 3* | 0.94955 | -0.31361 | 0.0625 | 0.215 |  | 627 | 2070 | 0.574 |
| *Ceratina sp. 4* | -0.10323 | -0.99466 | 0.0045 | 0.912 |  | 659 | 1563 | 0.066 |
| *Ceratina sp. 5* | 0.84049 | -0.54183 | 0.0125 | 0.756 |  | 669 | 1633 | 0.024 |
| *Ceratina sp. 6* | -0.92039 | -0.39101 | 0.009 | 0.819 |  | 669 | 2135 | 0.114 |
| *Ceratina sp. 7* | -0.30357 | -0.95281 | 0.0512 | 0.291 |  | 659 | 1608 | 0.197 |
| *Ceratina sp. 8* | -0.32622 | -0.94529 | 0.088 | 0.097 | . | 529 | 1563 | 0.066 |
| *Ceratina sp. 9* | -0.44305 | -0.8965 | 0.017 | 0.693 |  | 671 | 1671 | 0.090 |
| *Ceratina sp. 10* | 0.95698 | -0.29014 | 0.0419 | 0.343 |  | 1404 | 1864 | 0.090 |
| *Ceratina sp. 11* | 0.27288 | 0.96205 | 0.0121 | 0.672 |  | 2043 | 2043 | 0.042 |
| ***Ceratina sp. 12*** | **-0.45534** | **-0.89032** | **0.1164** | **0.048** | ***** | **671** | **1632** | 0.335 |
| *Ceratina sp. 12a* | -0.98757 | -0.1572 | 0.0276 | 0.478 |  | 1053 | 1528 | 0.012 |
| *Ceratina sp. 13* | 0.92277 | 0.38535 | 0.0289 | 0.518 |  | 1530 | 2192 | 0.024 |
| *Ceratina sp. 15* | 0.67931 | 0.73385 | 0.0298 | 0.521 |  | 1530 | 2070 | 0.030 |
| *Ceratina sp. 16* | 0.53255 | -0.8464 | 0.0255 | 0.433 |  | 1563 | 1563 | 0.006 |
| *Ceylalictus sp. 1* | -0.51879 | -0.8549 | 0.0837 | 0.132 |  | 627 | 1344 | 0.042 |
| *Cleptoparasitic sp.* | -0.98306 | -0.18328 | 0.0225 | 0.442 |  | 895 | 895 | 0.012 |
| *Coelioxys sp. 1* | 0.22682 | -0.97394 | 0.0296 | 0.495 |  | 730 | 2528 | 0.066 |
| *Coelioxys sp. 2* | 0.50711 | -0.86188 | 0.0083 | 0.792 |  | 1344 | 1344 | 0.006 |
| *Colletes sp. 1* | 0.33721 | -0.94143 | 0.0394 | 0.39 |  | 627 | 2043 | 0.054 |
| ***Colletes sp. 2*** | **-0.31812** | **-0.94805** | **0.1179** | **0.05** | ***** | **627** | **2528** | 0.048 |
| ***Crocisaspidia sp. 1*** | **-0.77694** | **-0.62958** | **0.1667** | **0.021** | ***** | **526** | **1630** | 0.102 |
| *Eupetersia sp. 1* | -0.52065 | -0.85377 | 0.0711 | 0.226 |  | 730 | 730 | 0.006 |
| *Eupetersia sp. 2* | 0.99899 | -0.0449 | 0.0258 | 0.409 |  | 2335 | 2335 | 0.006 |
| *Heriades sp. 1* | 0.62084 | -0.78394 | 0.063 | 0.223 |  | 730 | 2387 | 0.383 |
| *Heriades sp. 2* | 0.51953 | -0.85445 | 0.0784 | 0.121 |  | 526 | 2387 | 0.646 |
| *Heriades sp. 3* | 0.26215 | -0.96503 | 0.0012 | 1 |  | 1218 | 1218 | 0.006 |
| *Hylaeus sp. 1* | 0.71152 | -0.70267 | 0.0745 | 0.17 |  | 659 | 2528 | 0.197 |
| *Hylaeus sp. 2* | -0.132 | -0.99125 | 0.0492 | 0.302 |  | 671 | 2390 | 0.024 |
| ***Hypotrigona sp. 1*** | **-0.52737** | **-0.84964** | **0.1768** | **0.016** | ***** | **627** | **1094** | 0.269 |
| ***Hypotrigona sp. 2*** | **-0.59158** | **-0.80625** | **0.1603** | **0.029** | ***** | **627** | **1053** | 0.018 |
| *Lasioglossum sp. 1* | 0.51718 | -0.85588 | 0.0686 | 0.194 |  | 526 | 2528 | 0.700 |
| ***Lasioglossum sp. 2*** | **0.87592** | **0.48245** | **0.1403** | **0.031** | ***** | **669** | **2528** | 0.999 |
| *Lasioglossum sp. 2a* | 0.47015 | 0.88259 | 0.0285 | 0.509 |  | 1544 | 1700 | 0.078 |
| *Lasioglossum sp. 3* | 0.81917 | 0.57355 | 0.0768 | 0.152 |  | 1528 | 2528 | 0.197 |
| *Lasioglossum sp. 4* | 0.91248 | 0.40912 | 0.0238 | 0.555 |  | 2043 | 2528 | 0.012 |
| *Lasioglossum sp. 5* | 0.99111 | 0.13304 | 0.0341 | 0.437 |  | 526 | 2528 | 0.167 |
| ***Lasioglossum sp. a*** | **0.9688** | **0.24785** | **0.1221** | **0.04** | ***** | **669** | **2528** | 0.353 |
| *Lasioglossum sp. b* | -0.85932 | -0.51143 | 0.0127 | 0.686 |  | 671 | 671 | 0.006 |
| *Leuconomia sp. 1* | 0.87496 | 0.4842 | 0.0094 | 0.799 |  | 671 | 2070 | 0.173 |
| *Leuconomia sp. 2* | -0.18461 | -0.98281 | 0.0323 | 0.457 |  | 577 | 1632 | 0.072 |
| *Lipotriches sp. 1* | 0.61519 | -0.78838 | 0.0162 | 0.688 |  | 669 | 1632 | 0.036 |
| *Lipotriches sp. 1a* | 0.03694 | 0.99932 | 0.0188 | 0.507 |  | 1633 | 1633 | 0.006 |
| ***Lipotriches sp. 2*** | **-0.41744** | **-0.90871** | **0.1434** | **0.033** | ***** | **679** | **1624** | 0.030 |
| ***Lipotriches sp. 4*** | **-0.58862** | **-0.80841** | **0.2304** | **0.004** | ****** | **669** | **1462** | 0.084 |
| *Lipotriches sp. 5* | 0.97539 | 0.22048 | 0.058 | 0.253 |  | 659 | 2192 | 0.066 |
| *Lipotriches sp. 6* | 0.54067 | 0.84123 | 0.0237 | 0.569 |  | 671 | 1633 | 0.221 |
| *Lipotriches sp. 8* | 0.2671 | -0.96367 | 0.0339 | 0.402 |  | 529 | 1608 | 0.203 |
| *Lipotriches sp. 12* | -0.01826 | -0.99983 | 0.0306 | 0.479 |  | 679 | 2043 | 0.072 |
| *Lipotriches sp. 13* | -0.94517 | -0.32658 | 0.0246 | 0.508 |  | 671 | 1544 | 0.066 |
| ***Lipotriches sp. 14*** | **-0.39976** | **-0.91662** | **0.1486** | **0.024** | ***** | **526** | **1544** | 0.096 |
| *Lipotriches sp. 15* | -0.51541 | -0.85694 | 0.0033 | 0.923 |  | 669 | 669 | 0.006 |
| *Lipotriches sp. 16* | -0.85932 | -0.51143 | 0.0127 | 0.686 |  | 671 | 671 | 0.006 |
| *Lipotriches sp. 17* | -0.89811 | 0.43976 | 0.0512 | 0.331 |  | 577 | 577 | 0.006 |
| *Lipotriches sp. 18* | -0.73406 | 0.67909 | 0.0082 | 0.796 |  | 895 | 1633 | 0.036 |
| *Lipotriches sp. a* | 0.78889 | 0.61454 | 0.0496 | 0.298 |  | 1539 | 2135 | 0.078 |
| *Lipotriches sp. B* | -0.53654 | -0.84387 | 0.1279 | 0.067 | . | 669 | 895 | 0.024 |
| ***Macrogalea candida*** | **-0.43168** | **-0.90203** | **0.1759** | **0.011** | ***** | **526** | **1563** | 0.335 |
| *Macrogalea sp. 2* | -0.91572 | -0.40181 | 0.0759 | 0.206 |  | 529 | 529 | 0.018 |
| *Macronomia sp. 1* | 0.91062 | 0.41323 | 0.0006 | 0.997 |  | 740 | 1563 | 0.012 |
| ***Macronomia sp. 2*** | **-0.49579** | **-0.86844** | **0.2252** | **0.009** | ****** | **526** | **1462** | 0.042 |
| *Megachile ferina* | -0.98433 | -0.17636 | 0.0435 | 0.324 |  | 529 | 1630 | 0.048 |
| *Megachile sp. 1* | 0.60594 | -0.79551 | 0.0479 | 0.322 |  | 659 | 2035 | 0.161 |
| *Megachile sp. 1a* | -0.32402 | -0.94605 | 0.0461 | 0.289 |  | 577 | 577 | 0.018 |
| *Megachile sp. 1b* | 0.44271 | 0.89667 | 0.0148 | 0.752 |  | 669 | 1218 | 0.012 |
| *Megachile sp. 1c* | -0.89811 | 0.43976 | 0.0512 | 0.331 |  | 1608 | 1633 | 0.006 |
| *Megachile sp. 2* | 0.71657 | 0.69752 | 0.011 | 0.796 |  | 1632 | 1633 | 0.012 |
| *Megachile sp. 3* | 0.54103 | -0.841 | 0.0443 | 0.371 |  | 659 | 659 | 0.012 |
| *Megachile sp. 4* | -0.23559 | -0.97185 | 0.0549 | 0.248 |  | 526 | 1695 | 0.102 |
| *Megachile sp. 4a* | -0.5377 | -0.84314 | 0.1263 | 0.093 | . | 679 | 679 | 0.006 |
| *Megachile sp. 5* | 0.09572 | 0.99541 | 0.0014 | 0.971 |  | 659 | 1695 | 0.084 |
| ***Megachile sp. 5a*** | **-0.97365** | **-0.22805** | **0.1435** | **0.032** | ***** | **627** | **1630** | 0.036 |
| *Megachile sp. 6* | -0.3519 | -0.93604 | 0.0416 | 0.382 |  | 526 | 1630 | 0.024 |
| *Megachile sp. 7* | 0.6298 | 0.77675 | 0.0103 | 0.77 |  | 627 | 2528 | 0.197 |
| ***Megachile sp. 9*** | **-0.30913** | **-0.95102** | **0.224** | **0.006** | ****** | **526** | **1630** | 0.036 |
| *Megachile sp. 10* | 0.06887 | -0.99763 | 0.0531 | 0.234 |  | 659 | 1624 | 0.036 |
| *Megachile sp. 11* | -0.15 | 0.98869 | 0.0883 | 0.18 |  | 1503 | 1503 | 0.006 |
| *Megachile sp. 12* | -0.77541 | -0.63146 | 0.0368 | 0.393 |  | 529 | 1218 | 0.018 |
| ***Megachile sp. 13*** | **-0.59054** | **-0.80701** | **0.1437** | **0.029** | ***** | **526** | **2528** | 0.108 |
| *Megachile sp. 14* | -0.8104 | -0.58587 | 0.0151 | 0.702 |  | 627 | 2528 | 0.102 |
| *Megachile sp. 14a* | -0.61591 | -0.78782 | 0.0748 | 0.173 |  | 679 | 740 | 0.036 |
| *Megachile sp. 15* | -0.08033 | 0.99677 | 0.0361 | 0.369 |  | 1671 | 1671 | 0.006 |
| ***Megachile sp. 16*** | **-0.95645** | **-0.29188** | **0.1206** | **0.048** | ***** | **529** | **1630** | 0.042 |
| *Megachile sp. 17* | -0.5377 | -0.84314 | 0.1263 | 0.093 | . | 679 | 679 | 0.012 |
| *Megachile sp. 18* | 0.91789 | -0.39684 | 0.0365 | 0.401 |  | 1344 | 2528 | 0.024 |
| *Megachile sp. 19* | 0.50711 | -0.86188 | 0.0083 | 0.792 |  | 1344 | 1344 | 0.006 |
| *Meliponula ferruginea* | -0.4982 | -0.86706 | 0.1295 | 0.057 | . | 529 | 1624 | 0.155 |
| *Nomada sp. 1* | 0.94469 | 0.32797 | 0.0162 | 0.606 |  | 2135 | 2135 | 0.006 |
| *Nubenomia reichardia* | -0.68432 | -0.72918 | 0.0415 | 0.327 |  | 529 | 1624 | 0.484 |
| *Pachymelus sp. 1* | -0.5377 | -0.84314 | 0.1263 | 0.093 | . | 679 | 679 | 0.012 |
| *Pachynomia sp. 1* | 0.8398 | -0.5429 | 0.0004 | 0.991 |  | 669 | 2043 | 0.114 |
| *Parasitic sp. 1* | -0.52065 | -0.85377 | 0.0711 | 0.226 |  | 730 | 730 | 0.006 |
| *Parasitic sp. 2* | 0.77939 | -0.62654 | 0.029 | 0.474 |  | 659 | 1530 | 0.012 |
| *Plebeina armata* | -0.98306 | -0.18328 | 0.0225 | 0.442 |  | 895 | 895 | 0.012 |
| ***Plebeina sp. 1*** | **-0.61042** | **-0.79207** | **0.2789** | **0.005** | ****** | **577** | **1094** | 0.179 |
| *Pseudapis sp. 1* | 0.34631 | -0.93812 | 0.0142 | 0.691 |  | 577 | 2387 | 0.167 |
| *Pseudapis sp. 1a* | -0.90463 | -0.4262 | 0.1059 | 0.07 | . | 529 | 1053 | 0.084 |
| *Pseudapis sp. 2* | -0.99464 | -0.10342 | 0.0184 | 0.619 |  | 577 | 669 | 0.018 |
| *Pseudapis sp. 3* | 0.99534 | 0.0964 | 0.0199 | 0.576 |  | 669 | 2387 | 0.191 |
| *Pseudoanthidium sp. 1* | 0.95758 | -0.28818 | 0.06 | 0.208 |  | 669 | 2528 | 0.203 |
| ***Pseudoanthidium sp. 2*** | **-0.91945** | **-0.3932** | **0.1826** | **0.006** | ****** | **529** | **1630** | 0.197 |
| *Pseudoanthidium sp. 3* | 0.60095 | -0.79928 | 0.0344 | 0.463 |  | 669 | 2070 | 0.030 |
| *Pseudoanthidium sp. 4* | -0.51541 | -0.85694 | 0.0033 | 0.923 |  | 669 | 669 | 0.012 |
| *Pseudoanthidium sp. 5* | -0.18552 | 0.98264 | 0.0086 | 0.785 |  | 1630 | 1630 | 0.006 |
| *Pseudoanthidium sp. 6* | -0.73855 | 0.6742 | 0.0186 | 0.548 |  | 916 | 916 | 0.006 |
| *Seladonia sp. 1* | 0.90936 | -0.41601 | 0.0987 | 0.092 | . | 577 | 2528 | 0.712 |
| *Seladonia sp. 2* | 0.98401 | 0.1781 | 0.016 | 0.694 |  | 730 | 2070 | 0.072 |
| *Seladonia sp. 5* | 0.27288 | 0.96205 | 0.0121 | 0.672 |  | 2043 | 2043 | 0.006 |
| ***Steganomus sp. 1*** | **-0.94279** | **-0.33338** | **0.1378** | **0.043** | ***** | **529** | **1053** | 0.066 |
| *Steganomus sp. 2* | 0.65564 | 0.75507 | 0.0052 | 0.846 |  | 1530 | 1530 | 0.006 |
| *Steganomus sp. 3* | -0.78697 | -0.61699 | 0.0906 | 0.097 | . | 671 | 1528 | 0.024 |
| *Tetralonia sp. 1* | -0.84684 | -0.53184 | 0.067 | 0.204 |  | 669 | 669 | 0.084 |
| *Tetralonia sp. 2* | -0.28783 | -0.95768 | 0.0395 | 0.35 |  | 730 | 1864 | 0.030 |
| *Tetralonia sp. 3* | -0.11602 | -0.99325 | 0.1001 | 0.077 | . | 1218 | 1218 | 0.048 |
| *Tetralonia sp. 4* | 0.26215 | -0.96503 | 0.0012 | 1 |  | 895 | 895 | 0.012 |
| *Tetralonia sp. 5* | -0.98306 | -0.18328 | 0.0225 | 0.442 |  | 526 | 1624 | 0.006 |
| *Thrinchostoma sp. 1* | -0.08033 | 0.99677 | 0.0361 | 0.369 |  | 1671 | 1671 | 0.006 |
| *Thyreus sp. 1* | -0.03663 | -0.99933 | 0.0734 | 0.183 |  | 669 | 2528 | 0.084 |
| ***Trinomis sp. 1*** | **-0.48208** | **-0.87613** | **0.2726** | **0.001** | ******* | **526** | **1624** | 0.155 |
| *Unknown sp. 1* | -0.54375 | -0.83925 | 0.0681 | 0.204 |  | 679 | 1695 | # |
| *Unknown sp. 2* | -0.91572 | -0.40181 | 0.0759 | 0.206 |  | 529 | 529 | # |
| *Xylocopa caffra* | 0.9412 | 0.33786 | 0.0188 | 0.618 |  | 627 | 2528 | 0.365 |
| ***Xylocopa calens*** | **-0.59649** | **-0.80262** | **0.2934** | **0.002** | ****** | **627** | **730** | 0.018 |
| ***Xylocopa flavicolis*** | **-0.20202** | **-0.97938** | **0.1644** | **0.025** | ***** | **981** | **1558** | 0.024 |
| *Xylocopa flavorufa* | 0.74859 | -0.66303 | 0.0728 | 0.171 |  | 981 | 1864 | 0.138 |
| *Xylocopa inconstans* | 0.99677 | -0.08034 | 0.0156 | 0.704 |  | 669 | 1632 | 0.030 |
| *Xylocopa nigrita* | 0.47613 | -0.87938 | 0.0131 | 0.707 |  | 1053 | 1864 | 0.084 |
| *Xylocopa sp. 1* | 0.99487 | -0.10117 | 0.0196 | 0.633 |  | 916 | 1831 | 0.257 |
| *Xylocopa sp. 2* | 0.56989 | -0.82172 | 0.0356 | 0.444 |  | 981 | 1864 | 0.108 |
| *Xylocopa sp. 3* | -0.21756 | -0.97605 | 0.0541 | 0.239 |  | 895 | 1624 | 0.048 |
| *Zonalictus sp. 1* | 0.677 | -0.73599 | 0.0519 | 0.288 |  | 529 | 2528 | 0.215 |
| *Zonalictus sp. 2* | 0.48796 | -0.87287 | 0.0657 | 0.197 |  | 740 | 2192 | 0.054 |
| *Zonalictus sp. 3* | 0.69345 | 0.7205 | 0.1246 | 0.052 | . | 577 | 2528 | 0.652 |
| *Zonalictus sp. 4* | 0.9709 | -0.23948 | 0.0959 | 0.091 | . | 1344 | 2528 | 0.096 |
| *Zonalictus sp. 5* | 0.9977 | 0.0678 | 0.0799 | 0.119 |  | 1218 | 2528 | 0.108 |
| *Zonalictus sp. 6* | 0.77162 | 0.63608 | 0.0645 | 0.198 |  | 1539 | 2192 | 0.144 |
| *Zonalictus sp. 7* | 0.94836 | 0.31719 | 0.0647 | 0.188 |  | 1671 | 2528 | 0.090 |
| *Zonalictus sp. 8* | 0.99924 | -0.03905 | 0.0449 | 0.316 |  | 730 | 2528 | 0.096 |
| *Zonalictus sp. 9* | -0.94045 | 0.33993 | 0.0416 | 0.373 |  | 577 | 2390 | 0.042 |

**Table 3:** Plant species list, the elevational distribution of plants and the percentage abundance per species across the two counties. Non-metric multidimensional scaling ordination distances (NMDS1 & NMDS 2), coefficient of variation (**R^2^**), significance in ordination spaces by intrinsic species (P-value & Sig.), Min_Ele (Minimum elevation of species occurrence), Max_Ele (Maximum elevation of species occurrence), yellow colours (Species occurrence in Taita only), green colours (Species occurrence in Muranga only), blue colours (Species occurrence in Taita and Muranga), % abun (percentage abundance of each species).

| **Plant species** | **NMDS1** | **NMDS2** | **R^2^** | **P-value** | **Sig.** | **Min_Ele** | **Max_Ele** | **% visit** |
| --- | --- | --- | --- | --- | --- | --- | --- | --- |
| *Abutilon fruticosum* | -0.31493 | 0.94911 | 0.0344 | 0.406 |  | 669 | 669 | 0.006 |
| *Abutilon guineense* | -0.72263 | 0.69123 | 0.0978 | 0.106 |  | 526 | 529 | 0.054 |
| *Abutilon longicuspe* | 0.99822 | -0.05967 | 0.0035 | 0.871 |  | 1671 | 1671 | 0.018 |
| *Abutilon mauritianum* | -0.50649 | 0.86225 | 0.0198 | 0.573 |  | 671 | 671 | 0.012 |
| *Abutilon sp.1* | -0.50649 | 0.86225 | 0.0198 | 0.573 |  | 671 | 671 | 0.018 |
| *Acacia bussei* | -0.71565 | 0.69846 | 0.0875 | 0.136 |  | 526 | 526 | 0.078 |
| *Acacia hockii* | 0.01959 | 0.99981 | 0.0498 | 0.278 |  | 1528 | 1528 | 0.024 |
| *Acacia senegalensis* | -0.63879 | 0.76938 | 0.0478 | 0.301 |  | 529 | 895 | 0.066 |
| *Acacia sp.1* | -0.83258 | 0.55391 | 0.0222 | 0.521 |  | 529 | 529 | 0.006 |
| *Acanthospermum glabratum* | 0.22514 | -0.97433 | 0.04 | 0.358 |  | 1563 | 2043 | 0.012 |
| *Acanthospermum hispidum* | -0.68951 | -0.72428 | 0.0521 | 0.279 |  | 529 | 1352 | 0.012 |
| *Achyranthes aspera* | 0.253 | -0.96747 | 0.0248 | 0.503 |  | 1608 | 2043 | 0.018 |
| *Achyranthes sp.1* | -0.67333 | -0.73934 | 0.0027 | 0.911 |  | 981 | 981 | 0.006 |
| *Achyrospermum schimperi* | 0.66806 | -0.7441 | 0.0576 | 0.231 |  | 2035 | 2192 | 0.311 |
| *Acmella caulirhiza* | 0.50905 | -0.86074 | 0.0651 | 0.165 |  | 2070 | 2070 | 0.030 |
| *Aeollanthus repens* | -0.09086 | -0.99586 | 0.0637 | 0.171 |  | 1831 | 1831 | 0.012 |
| *Aeschynomene schimperi* | 0.01959 | 0.99981 | 0.0498 | 0.278 |  | 1528 | 1528 | 0.084 |
| *Agauria salicifolia* | -0.33829 | -0.94104 | 0.0604 | 0.208 |  | 1352 | 1352 | 0.012 |
| *Ageratum conyzoides* | 0.56131 | -0.8276 | 0.0642 | 0.196 |  | 1344 | 2390 | 0.221 |
| *Ajuga integrifolia* | 0.14243 | 0.9898 | 0.0786 | 0.144 |  | 1462 | 1576 | 1.441 |
| *Aloe myriacantha* | -0.27906 | 0.96027 | 0.0008 | 0.983 |  | 1624 | 1624 | 0.006 |
| *Aloe secundiflora* | -0.5266 | 0.85011 | 0.0266 | 0.479 |  | 671 | 740 | 0.132 |
| *Alternanthera sessilis* | -0.83258 | 0.55391 | 0.0222 | 0.521 |  | 529 | 529 | 0.012 |
| *Amaranthus dubius* | -0.02682 | -0.99964 | 0.0002 | 0.988 |  | 1576 | 1633 | 0.024 |
| *Aneilemia aequinoctiale* | 0.05461 | -0.99851 | 0.0521 | 0.277 |  | 1864 | 1864 | 0.598 |
| *Argyrolobium fischeri* | 0.8811 | 0.47293 | 0.0423 | 0.381 |  | 2528 | 2528 | 0.155 |
| *Aristida adscensionis* | -0.6574 | 0.75355 | 0.0533 | 0.262 |  | 740 | 740 | 0.018 |
| *Aspilia mossambicensis* | 0.0599 | -0.9982 | 0.008 | 0.762 |  | 529 | 1624 | 1.232 |
| *Asystasia mysorensis* | 0.21621 | -0.97635 | 0.0117 | 0.74 |  | 1608 | 1608 | 0.042 |
| *Barleria taitensis* | -0.99953 | 0.03054 | 0.0376 | 0.379 |  | 627 | 627 | 0.006 |
| *Bidens pilosa* | 0.19748 | -0.98031 | 0.0742 | 0.165 |  | 669 | 2414 | 4.562 |
| *Bidens sp.1* | -0.09086 | -0.99586 | 0.0637 | 0.171 |  | 1831 | 1831 | 0.598 |
| *Blepharis edulis* | -0.42554 | 0.90494 | 0.0965 | 0.081 | . | 526 | 679 | 0.377 |
| *Bothriochloa insculpta* | -0.09512 | -0.99547 | 0.0067 | 0.839 |  | 671 | 1563 | 0.024 |
| *Brachiaria sp.1* | 0.18792 | -0.98218 | 0.0122 | 0.695 |  | 1558 | 1632 | 0.066 |
| *Buddleja polystachya* | 0.31489 | -0.94913 | 0.0181 | 0.614 |  | 2043 | 2043 | 0.072 |
| *Caesalpinia decapetala* | 0.01488 | -0.99989 | 0.0323 | 0.454 |  | 1218 | 1632 | 0.419 |
| *Calliandra houstoniana* | 0.05905 | -0.99826 | 0.1258 | 0.056 | . | 1344 | 1762 | 0.245 |
| *Callistemon viminalis* | 0.21621 | -0.97635 | 0.0117 | 0.74 |  | 1608 | 1608 | 0.108 |
| *Calotropis procera* | -0.70066 | 0.7135 | 0.0334 | 0.406 |  | 529 | 669 | 0.030 |
| *Caylusea abyssinica* | 0.39289 | -0.91959 | 0.002 | 0.945 |  | 1695 | 1695 | 0.006 |
| *Chamaecrista mimosoides* | -0.05699 | -0.99837 | 0.0277 | 0.516 |  | 1344 | 1344 | 0.006 |
| *Chamaecrista sp.1* | -0.53972 | -0.84185 | 0.0033 | 0.921 |  | 1218 | 1624 | 0.048 |
| *Cissus aralioides* | -0.45074 | 0.89266 | 0.0285 | 0.481 |  | 895 | 895 | 0.018 |
| *Clerodendrum eriophyllum* | -0.40866 | 0.91269 | 0.0009 | 0.966 |  | 730 | 1624 | 0.126 |
| *Clinopodium abyssinicum* | 0.92101 | 0.38953 | 0.0437 | 0.335 |  | 2043 | 2528 | 0.072 |
| *Commelina benghalensis* | -0.45074 | 0.89266 | 0.0285 | 0.481 |  | 895 | 2414 | 0.036 |
| *Commelina sp.1* | -0.31493 | 0.94911 | 0.0344 | 0.406 |  | 669 | 669 | 0.096 |
| *Convolvulus kilimandschari* | 0.91754 | -0.39764 | 0.0776 | 0.159 |  | 2390 | 2390 | 0.006 |
| *Conyza bonariensis* | 0.60722 | -0.79454 | 0.0687 | 0.193 |  | 1352 | 2528 | 0.090 |
| *Conyza newii* | 0.05461 | -0.99851 | 0.0521 | 0.277 |  | 1864 | 1864 | 1.244 |
| *Conyza schimperi* | 0.79656 | -0.60456 | 0.0315 | 0.424 |  | 2192 | 2192 | 0.006 |
| *Conyza steudelii* | 0.56888 | -0.82242 | 0.0621 | 0.209 |  | 2135 | 2192 | 0.012 |
| *Cordia africana* | 0.23556 | -0.97186 | 0.0068 | 0.802 |  | 1632 | 1632 | 0.006 |
| *Crambe cordifolia* | -0.05699 | -0.99837 | 0.0277 | 0.516 |  | 1344 | 1344 | 0.006 |
| *Crassocephalum montuosum* | 0.43362 | -0.9011 | 0.0532 | 0.247 |  | 2043 | 2192 | 0.688 |
| *Crassocephalum picridifolium* | 0.39289 | -0.91959 | 0.002 | 0.945 |  | 1695 | 1695 | 0.012 |
| *Crassocephalum vitellinum* | 0.72679 | -0.68686 | 0.052 | 0.24 |  | 2043 | 2528 | 0.377 |
| *Crotalaria axillaris* | -0.05699 | -0.99837 | 0.0277 | 0.516 |  | 1344 | 1344 | 0.036 |
| *Crotalaria barkae* | -0.33895 | 0.9408 | 0.1519 | 0.034 | * | 627 | 659 | 0.054 |
| *Crotalaria ukambensis* | -0.61161 | 0.79116 | 0.0423 | 0.312 |  | 679 | 679 | 0.006 |
| *Croton bonplandianus* | -0.67333 | -0.73934 | 0.0027 | 0.911 |  | 981 | 981 | 0.018 |
| *Cucumis dipsaceus* | -0.825 | -0.56514 | 0.0073 | 0.76 |  | 1053 | 1053 | 0.006 |
| *Cyathula cylindrica* | 0.07794 | -0.99696 | 0.0514 | 0.263 |  | 1864 | 2528 | 0.634 |
| *cyathula orthacantha* | -0.31493 | 0.94911 | 0.0344 | 0.406 |  | 669 | 669 | 0.006 |
| *Cynodon nlemfuensis* | -0.1312 | -0.99136 | 0.0022 | 0.946 |  | 1576 | 1633 | 0.114 |
| *Cyperus tomaiophyllus* | 0.8811 | 0.47293 | 0.0423 | 0.381 |  | 2528 | 2528 | 0.024 |
| *Cyphostemma kilimandscharicum* | 0.8811 | 0.47293 | 0.0423 | 0.381 |  | 2528 | 2528 | 0.060 |
| *Dactyloctenium aegyptium* | -0.50649 | 0.86225 | 0.0198 | 0.573 |  | 671 | 671 | 0.012 |
| *Datura suaveolens* | 0.50905 | -0.86074 | 0.0651 | 0.165 |  | 2070 | 2070 | 0.359 |
| *Desmodium incanum* | -0.33829 | -0.94104 | 0.0604 | 0.208 |  | 1352 | 1352 | 0.006 |
| *Desmodium intortum* | -0.18021 | -0.98363 | 0.0469 | 0.297 |  | 1344 | 1404 | 0.018 |
| *Desmodium sp.1* | -0.37431 | -0.92731 | 0.0778 | 0.141 |  | 529 | 1352 | 0.036 |
| *Desmodium uncinatum* | 0.70849 | -0.70572 | 0.0083 | 0.773 |  | 1558 | 1630 | 0.090 |
| *Dicliptera paniculata* | -0.87784 | -0.47895 | 0.0158 | 0.624 |  | 730 | 730 | 0.012 |
| *Digera muricata* | -0.45162 | 0.89221 | 0.2659 | 0.001 | *** | 526 | 895 | 0.921 |
| *Dyschoriste clinopodioides* | 0.79656 | -0.60456 | 0.0315 | 0.424 |  | 2192 | 2192 | 0.012 |
| *Dyschoriste hildebrandtii* | 0.21089 | 0.97751 | 0.0149 | 0.679 |  | 1462 | 1462 | 0.018 |
| *Emilia discifolia* | 0.28557 | 0.95836 | 0.0099 | 0.789 |  | 1503 | 1695 | 0.054 |
| *Entada leptostachya* | 0.21089 | 0.97751 | 0.0149 | 0.679 |  | 1462 | 1462 | 0.066 |
| *Eragrostis superba* | -0.46353 | 0.88608 | 0.0149 | 0.663 |  | 577 | 1624 | 0.096 |
| *Eriobotrya japonica* | 0.54165 | -0.8406 | 0.0628 | 0.205 |  | 2135 | 2390 | 0.024 |
| *Eriosema sp.1* | 0.8811 | 0.47293 | 0.0423 | 0.381 |  | 2528 | 2528 | 0.006 |
| *Euclea racemosa* | -0.31599 | -0.94876 | 0.008 | 0.739 |  | 1218 | 1218 | 0.006 |
| *Euphorbia bicompacta* | -0.45074 | 0.89266 | 0.0285 | 0.481 |  | 895 | 895 | 0.012 |
| *Euphorbia cuneata* | -0.99828 | -0.05866 | 0.048 | 0.289 |  | 627 | 1094 | 0.161 |
| *Euphorbia heterophylla* | 0.21621 | -0.97635 | 0.0117 | 0.74 |  | 1608 | 1608 | 0.030 |
| *Euphorbia hirta* | -0.31493 | 0.94911 | 0.0344 | 0.406 |  | 669 | 669 | 0.006 |
| *Euryops chrysanthemoides* | 0.54848 | -0.83616 | 0.0629 | 0.209 |  | 981 | 2414 | 0.945 |
| *Galinsoga parviflora* | 0.15904 | -0.98727 | 0.0278 | 0.47 |  | 1563 | 1632 | 0.155 |
| *Galinsoga quadriradiata* | 0.504 | -0.8637 | 0.028 | 0.472 |  | 1633 | 2192 | 0.173 |
| *Geranium vagans* | 0.8811 | 0.47293 | 0.0423 | 0.381 |  | 2528 | 2528 | 0.030 |
| *Grevillea robusta* | 0.49783 | 0.86727 | 0.0001 | 1 |  | 1633 | 1633 | 0.006 |
| *Grewia kakothamnos* | -0.99953 | 0.03054 | 0.0376 | 0.379 |  | 627 | 627 | 0.006 |
| *Grewia tephrodermis* | -0.50272 | 0.86445 | 0.1359 | 0.044 | * | 526 | 1352 | 0.592 |
| *Grewia villosa* | -0.51808 | 0.85533 | 0.0013 | 0.975 |  | 895 | 1695 | 0.018 |
| *Gutenbergia cordifolia* | -0.02308 | -0.99973 | 0.0631 | 0.224 |  | 730 | 1762 | 1.196 |
| *Harpachne schimperi* | -0.27906 | 0.96027 | 0.0008 | 0.983 |  | 1624 | 1624 | 0.012 |
| *Helichrysum forskahlii* | 0.06727 | -0.99774 | 0.0196 | 0.604 |  | 1558 | 1558 | 0.006 |
| *Helichrysum glumaceum* | -0.6574 | 0.75355 | 0.0533 | 0.262 |  | 740 | 740 | 0.006 |
| *Helichrysum odoratissimum* | -0.33829 | -0.94104 | 0.0604 | 0.208 |  | 1352 | 1352 | 0.132 |
| *Helichrysum schimperi* | 0.05461 | -0.99851 | 0.0521 | 0.277 |  | 1864 | 1864 | 0.622 |
| *Heliotropium steudneri* | -0.48037 | 0.87706 | 0.0334 | 0.418 |  | 529 | 895 | 0.060 |
| *Hermannia exappendiculata* | -0.61161 | 0.79116 | 0.0423 | 0.312 |  | 679 | 679 | 0.084 |
| *Hermannia oliveri* | -0.61161 | 0.79116 | 0.0423 | 0.312 |  | 679 | 679 | 0.006 |
| *Hewittia malabarica* | 0.06727 | -0.99774 | 0.0196 | 0.604 |  | 1558 | 1558 | 0.018 |
| *Hibiscus acetosella* | 0.48968 | -0.8719 | 0.0154 | 0.662 |  | 2035 | 2035 | 0.006 |
| *Hibiscus kabuyeana* | -0.87784 | -0.47895 | 0.0158 | 0.624 |  | 730 | 730 | 0.006 |
| *Hibiscus micranthus* | -0.6574 | 0.75355 | 0.0533 | 0.262 |  | 740 | 740 | 0.006 |
| *Hibiscus sidiformis* | -0.31493 | 0.94911 | 0.0344 | 0.406 |  | 669 | 669 | 0.006 |
| *Hypericum revolutum* | 0.8811 | 0.47293 | 0.0423 | 0.381 |  | 2528 | 2528 | 0.048 |
| *Hypoestes forskaolii* | 0.21089 | 0.97751 | 0.0149 | 0.679 |  | 1462 | 1462 | 0.012 |
| *Hypoxis obtusa* | 0.27807 | 0.96056 | 0.0013 | 0.966 |  | 1563 | 1563 | 0.012 |
| *Hyptis suaveolens* | 0.06727 | -0.99774 | 0.0196 | 0.604 |  | 1558 | 1558 | 0.006 |
| *Idiospermum kilimandscharica* | -0.87784 | -0.47895 | 0.0158 | 0.624 |  | 730 | 730 | 0.018 |
| *Impatiens hoehnelii* | 0.64969 | 0.7602 | 0.2878 | 0.017 | * | 2335 | 2335 | 0.042 |
| *Indigofera arrecta* | 0.19068 | -0.98165 | 0.0225 | 0.575 |  | 1558 | 1632 | 0.048 |
| *Indigofera colutea* | -0.35532 | 0.93474 | 0.0587 | 0.225 |  | 669 | 895 | 0.018 |
| *Indigofera schimperi* | -0.51052 | 0.85987 | 0.0402 | 0.336 |  | 577 | 671 | 0.054 |
| *Indigofera spicata* | 0.14253 | 0.98979 | 0.0126 | 0.715 |  | 1539 | 1633 | 0.018 |
| *Indigofera tinctoria* | 0.09951 | 0.99504 | 0.0148 | 0.655 |  | 529 | 1462 | 0.042 |
| *Indigofera vohemarensis* | -0.31599 | -0.94876 | 0.008 | 0.739 |  | 1218 | 1218 | 0.072 |
| *Ipomoea batatas* | 0.97943 | -0.20177 | 0.0062 | 0.825 |  | 1630 | 1630 | 0.018 |
| *Ipomoea mombassana* | -0.87784 | -0.47895 | 0.0158 | 0.624 |  | 730 | 730 | 0.006 |
| *Ipomoea sp.1* | 0.16683 | -0.98599 | 0.0333 | 0.429 |  | 1563 | 1608 | 0.018 |
| *Justicia calyculata* | 0.25812 | -0.96611 | 0.0094 | 0.758 |  | 1462 | 2192 | 2.129 |
| *Justicia debilis* | -0.4887 | -0.87245 | 0.0167 | 0.646 |  | 730 | 1624 | 0.030 |
| *Justicia diclipteroides* | -0.88036 | -0.4743 | 0.0068 | 0.878 |  | 730 | 1633 | 0.024 |
| *Justicia flava* | -0.43778 | 0.89908 | 0.1513 | 0.039 | * | 577 | 895 | 0.036 |
| *Justicia striata* | 0.59263 | -0.80548 | 0.0038 | 0.895 |  | 916 | 2528 | 1.028 |
| *Kalanchoe lateritia* | -0.54146 | -0.84073 | 0.014 | 0.69 |  | 1053 | 1218 | 0.012 |
| *Kleinia squarrosa* | -0.45074 | 0.89266 | 0.0285 | 0.481 |  | 895 | 895 | 0.012 |
| *Kyllinga brevifolia* | 0.23547 | -0.97188 | 0.0136 | 0.692 |  | 1563 | 1632 | 2.308 |
| *Kyllinga bulbosa* | 0.44399 | -0.89603 | 0.0368 | 0.395 |  | 1558 | 2528 | 0.054 |
| *Kyllinga sp.1* | 0.79656 | -0.60456 | 0.0315 | 0.424 |  | 2192 | 2192 | 0.012 |
| *Lantana camara* | 0.25853 | -0.966 | 0.0031 | 0.926 |  | 981 | 1700 | 0.269 |
| *Leucaena leucocephala* | -0.67333 | -0.73934 | 0.0027 | 0.911 |  | 981 | 981 | 0.012 |
| *Leucas glabrata* | -0.97712 | -0.21268 | 0.0216 | 0.538 |  | 679 | 730 | 0.090 |
| *Leucas grandis* | 0.00539 | -0.99999 | 0.0789 | 0.131 |  | 529 | 1864 | 2.224 |
| *Leucas neuflizeana* | -0.90293 | 0.42978 | 0.0957 | 0.09 | . | 627 | 1053 | 0.102 |
| *Lippia javanica* | 0.91812 | -0.39629 | 0.0777 | 0.148 |  | 1462 | 2390 | 5.991 |
| *Lippia kituiensis* | -0.09616 | -0.99537 | 0.0294 | 0.452 |  | 1053 | 1344 | 0.084 |
| *Lobelia fervens* | 0.67788 | 0.73517 | 0.1252 | 0.073 | . | 1633 | 2528 | 0.018 |
| *Maerua endlichii* | -0.31493 | 0.94911 | 0.0344 | 0.406 |  | 669 | 669 | 0.006 |
| ***Maerua sp.1*** | **-0.26611** | **0.96394** | **0.1449** | **0.047** | ***** | **659** | **1404** | 6.494 |
| *Maerua triphylla* | -0.57289 | 0.81963 | 0.0949 | 0.103 |  | 669 | 895 | 0.514 |
| *Melanthera scandens* | 0.06727 | -0.99774 | 0.0196 | 0.604 |  | 1558 | 1558 | 0.006 |
| *Melhania ovata* | -0.51207 | 0.85895 | 0.039 | 0.351 |  | 577 | 895 | 0.138 |
| *Melinis repens* | 0.37848 | -0.92561 | 0.0338 | 0.437 |  | 2135 | 2135 | 0.006 |
| *Mimosa pudica* | 0.06727 | -0.99774 | 0.0196 | 0.604 |  | 1558 | 1558 | 0.006 |
| *Moligono sp.1* | 0.79656 | -0.60456 | 0.0315 | 0.424 |  | 2192 | 2192 | 0.006 |
| *Murdannia simplex* | -0.67333 | -0.73934 | 0.0027 | 0.911 |  | 981 | 981 | 0.018 |
| *Neonotonia wightii* | 0.26886 | 0.96318 | 0.0382 | 0.402 |  | 1053 | 1633 | 0.347 |
| *Nicandra physaloides* | 0.12525 | 0.99212 | 0.0564 | 0.203 |  | 1539 | 1539 | 0.012 |
| ***Ocimum americanum*** | **-0.67761** | **0.73542** | **0.2355** | **0.002** | ****** | **529** | **1624** | 0.305 |
| *Ocimum gratissimum* | 0.07903 | -0.99687 | 0.0851 | 0.12 |  | 1218 | 2043 | 7.468 |
| *Ocimum Kenyense* | 0.70942 | -0.70479 | 0.0459 | 0.295 |  | 2035 | 2192 | 0.341 |
| *Ocimum kilimandscharicum* | 0.25081 | -0.96804 | 0.0065 | 0.813 |  | 1563 | 1632 | 0.622 |
| *Ocimum sp.1* | -0.12384 | -0.9923 | 0.0336 | 0.418 |  | 981 | 1695 | 1.866 |
| *Oldenlandia friesiorum* | 0.8811 | 0.47293 | 0.0423 | 0.381 |  | 2528 | 2528 | 0.239 |
| *Oreosyce africana* | 0.97943 | -0.20177 | 0.0062 | 0.825 |  | 1630 | 1630 | 0.042 |
| *Ornithogalum tenuifolium* | 0.06928 | -0.9976 | 0.0079 | 0.775 |  | 1544 | 1544 | 0.006 |
| *Osteospermum vaillantii* | 0.44847 | 0.8938 | 0.0062 | 0.849 |  | 1530 | 1530 | 0.006 |
| *Oxalis corniculata* | 0.01216 | 0.99993 | 0.0704 | 0.173 |  | 1503 | 1563 | 0.060 |
| *Oxygonum sinuatum* | -0.97791 | -0.20904 | 0.0138 | 0.68 |  | 529 | 1633 | 0.245 |
| *Paederia pospischilii* | -0.99022 | -0.13954 | 0.0234 | 0.513 |  | 679 | 730 | 0.036 |
| *Panicum maximum* | -0.0065 | -0.99998 | 0.0115 | 0.727 |  | 1544 | 1633 | 0.149 |
| *Panicum trichocladum* | 0.39548 | 0.91848 | 0.0094 | 0.752 |  | 1462 | 1530 | 0.084 |
| ***Parochetus communis*** | **0.64969** | **0.7602** | **0.2878** | **0.017** | ***** | **2335** | **2335** | 0.072 |
| *Parthenium hysterophorus* | -0.15608 | -0.98774 | 0.0039 | 0.841 |  | 1576 | 1576 | 0.550 |
| *Passiflora subpeltata* | -0.26413 | 0.96449 | 0.1396 | 0.059 | . | 659 | 659 | 0.006 |
| *Pavonia burchellii* | 0.05461 | -0.99851 | 0.0521 | 0.277 |  | 1864 | 1864 | 0.018 |
| *Pavonia urens* | 0.50905 | -0.86074 | 0.0651 | 0.165 |  | 2070 | 2070 | 0.006 |
| *Pedaliodiscus macrocarpus* | -0.64287 | 0.76598 | 0.0223 | 0.527 |  | 669 | 1053 | 0.066 |
| *Pentas parvifolia* | -0.67333 | -0.73934 | 0.0027 | 0.911 |  | 981 | 981 | 0.006 |
| *Pentas zanzibarica* | 0.06195 | -0.99808 | 0.0385 | 0.383 |  | 1218 | 1864 | 0.144 |
| *Persea sp.1* | 0.25734 | -0.96632 | 0.1049 | 0.087 | . | 1558 | 2135 | 2.392 |
| *Persicaria senegalensis* | 0.51487 | -0.85727 | 0.0675 | 0.16 |  | 1462 | 2070 | 0.120 |
| *Persicaria setosula* | 0.50905 | -0.86074 | 0.0651 | 0.165 |  | 2070 | 2070 | 0.006 |
| *Phyllanthus sp.1* | 0.37848 | -0.92561 | 0.0338 | 0.437 |  | 2135 | 2135 | 0.006 |
| *Pittosporum viridiflorum* | -0.31599 | -0.94876 | 0.008 | 0.739 |  | 1218 | 1218 | 0.012 |
| *Plant sp. 1* | 0.8811 | 0.47293 | 0.0423 | 0.381 |  | 2528 | 2528 | 0.024 |
| *Plant sp. 2* | -0.0889 | -0.99604 | 0.0635 | 0.167 |  | 1831 | 2528 | 1.202 |
| *Plant sp. 3* | -0.05699 | -0.99837 | 0.0277 | 0.516 |  | 1344 | 1344 | 0.114 |
| *Plant sp. 4* | -0.05699 | -0.99837 | 0.0277 | 0.516 |  | 1344 | 1344 | 0.006 |
| *Plant sp. 5* | -0.05699 | -0.99837 | 0.0277 | 0.516 |  | 1344 | 1344 | 0.012 |
| *Plant sp. 6* | -0.31493 | 0.94911 | 0.0344 | 0.406 |  | 669 | 669 | 0.018 |
| *Plant sp. 7* | -0.51911 | 0.85471 | 0.0542 | 0.231 |  | 577 | 577 | 0.006 |
| *Plant sp. 8* | -0.45667 | 0.88964 | 0.0238 | 0.525 |  | 659 | 916 | 0.263 |
| *Plant sp. 9* | 0.01959 | 0.99981 | 0.0498 | 0.278 |  | 1528 | 1528 | 0.018 |
| *Plant sp. 10* | 0.06727 | -0.99774 | 0.0196 | 0.604 |  | 1558 | 1558 | 0.006 |
| *Plant sp. 11* | 0.46827 | 0.88358 | 0.0436 | 0.327 |  | 2414 | 2414 | 0.006 |
| *Plant sp. 12* | 0.06727 | -0.99774 | 0.0196 | 0.604 |  | 1558 | 1558 | 0.066 |
| ***Plant sp. 13*** | **0.57742** | **0.81645** | **0.1571** | **0.041** | ***** | **2387** | **2387** | 0.012 |
| *Plant sp. 14* | 0.51791 | -0.85543 | 0.0033 | 0.91 |  | 1700 | 1700 | 0.012 |
| *Plant sp. 15* | 0.58888 | 0.80822 | 0.2203 | 0.023 |  | 2335 | 2387 | 0.048 |
| *Plant sp. 16* | 0.51791 | -0.85543 | 0.0033 | 0.91 |  | 1700 | 1700 | 0.012 |
| ***Plant sp. 17*** | **0.63595** | **0.77173** | **0.1768** | **0.024** | ***** | **2387** | **2390** | 0.048 |
| *Plant sp. 18* | 0.06727 | -0.99774 | 0.0196 | 0.604 |  | 1558 | 1558 | 0.006 |
| ***Plant sp. 19*** | **0.57742** | **0.81645** | **0.1571** | **0.041** | ***** | **2387** | **2387** | 0.096 |
| *Plant sp. 20* | 0.46827 | 0.88358 | 0.0436 | 0.327 |  | 2414 | 2414 | 0.066 |
| *Plant sp. 21* | 0.46827 | 0.88358 | 0.0436 | 0.327 |  | 2414 | 2414 | 0.006 |
| ***Plant sp. 22*** | **0.57742** | **0.81645** | **0.1571** | **0.041** | ***** | **2387** | **2387** | 0.012 |
| *Plant sp. 23* | 0.8811 | 0.47293 | 0.0423 | 0.381 |  | 2528 | 2528 | 0.024 |
| *Plant sp. 24* | 0.91894 | 0.3944 | 0.0519 | 0.283 |  | 2390 | 2528 | 0.072 |
| *Plant sp. 25* | 0.06727 | -0.99774 | 0.0196 | 0.604 |  | 1558 | 1558 | 0.120 |
| *Plant sp. 26* | 0.99395 | 0.10981 | 0.0962 | 0.11 |  | 2390 | 2528 | 0.066 |
| ***Plant sp. 27*** | **0.57742** | **0.81645** | **0.1571** | **0.041** | ***** | **2387** | **2387** | 0.006 |
| *Plant sp. 28* | 0.06727 | -0.99774 | 0.0196 | 0.604 |  | 1558 | 1558 | 0.006 |
| *Plant sp. 29* | 0.06727 | -0.99774 | 0.0196 | 0.604 |  | 1558 | 1558 | 0.197 |
| ***Plant sp. 30*** | **0.64969** | **0.7602** | **0.2878** | **0.017** | ***** | **2335** | **2335** | 0.018 |
| ***Plant sp. 31*** | **0.3799** | **0.92503** | **0.2059** | **0.015** | ***** | **1528** | **2414** | 0.036 |
| *Plant sp. 32* | 0.51791 | -0.85543 | 0.0033 | 0.91 |  | 1700 | 1700 | 0.060 |
| *Plant sp. 33* | 0.51791 | -0.85543 | 0.0033 | 0.91 |  | 1700 | 1700 | 0.030 |
| *Plant sp. 34* | 0.91754 | -0.39764 | 0.0776 | 0.159 |  | 2390 | 2390 | 0.006 |
| *Plant sp. 35* | 0.97943 | -0.20177 | 0.0062 | 0.825 |  | 1630 | 1630 | 0.018 |
| *Platostoma africana* | 0.46681 | -0.88436 | 0.0511 | 0.26 |  | 2135 | 2414 | 0.024 |
| *Platostoma denticulatum* | 0.94496 | -0.3272 | 0.0619 | 0.221 |  | 2043 | 2528 | 0.239 |
| *Plectranthus alboviolaceus* | 0.99822 | -0.05967 | 0.0035 | 0.871 |  | 1671 | 1671 | 0.006 |
| *Plectranthus alpinus* | 0.93864 | -0.34491 | 0.0876 | 0.132 |  | 2390 | 2528 | 0.066 |
| *Plectranthus barbatus* | 0.97563 | -0.21942 | 0.0055 | 0.851 |  | 1671 | 2414 | 0.305 |
| *Plectranthus caninus* | 0.60623 | -0.79529 | 0.0139 | 0.654 |  | 1462 | 2035 | 0.090 |
| *Plectranthus ignarius* | -0.27906 | 0.96027 | 0.0008 | 0.983 |  | 1624 | 1624 | 0.036 |
| *Plectranthus kamerunensis* | -0.05935 | -0.99824 | 0.0631 | 0.181 |  | 1671 | 1831 | 0.048 |
| ***Plectranthus luteus*** | **0.64969** | **0.7602** | **0.2878** | **0.017** | ***** | **2335** | **2335** | 0.102 |
| *Plectranthus olostegioides* | -0.6574 | 0.75355 | 0.0533 | 0.262 |  | 740 | 740 | 0.006 |
| *Plectranthus punctatus* | 0.95714 | -0.28964 | 0.0761 | 0.169 |  | 2035 | 2528 | 1.315 |
| *Plectranthus sp.1* | -0.67333 | -0.73934 | 0.0027 | 0.911 |  | 981 | 981 | 0.006 |
| *Polygala sphenoptera* | -0.79972 | 0.60037 | 0.0627 | 0.223 |  | 730 | 740 | 0.018 |
| *Premna oligocephala* | -0.87784 | -0.47895 | 0.0158 | 0.624 |  | 730 | 730 | 0.006 |
| *Psidium guajava* | -0.05699 | -0.99837 | 0.0277 | 0.516 |  | 1344 | 1344 | 0.036 |
| *Psychotria capensis* | -0.31599 | -0.94876 | 0.008 | 0.739 |  | 1218 | 1218 | 0.640 |
| ***Psychotria mahonii*** | **0.61492** | **0.78859** | **0.3004** | **0.01** | ****** | **2335** | **2528** | 17.400 |
| *Pteridium aquilinum* | 0.8811 | 0.47293 | 0.0423 | 0.381 |  | 2528 | 2528 | 0.006 |
| *Pupalia lappacea* | -0.50649 | 0.86225 | 0.0198 | 0.573 |  | 671 | 671 | 0.012 |
| *Pycnostachys meyeri* | 0.91754 | -0.39764 | 0.0776 | 0.159 |  | 2390 | 2390 | 0.084 |
| *Pycnostachys sp.1* | 0.01959 | 0.99981 | 0.0498 | 0.278 |  | 1528 | 1528 | 0.006 |
| *Ranunculus multifidus* | 0.31489 | -0.94913 | 0.0181 | 0.614 |  | 2043 | 2043 | 0.006 |
| *Rhus natalensis* | 0.97943 | -0.20177 | 0.0062 | 0.825 |  | 1630 | 1630 | 0.108 |
| *Rhynchosia elegans* | 0.49783 | 0.86727 | 0.0001 | 1 |  | 1633 | 1633 | 0.006 |
| *Rhynchosia sp.1* | -0.51698 | 0.856 | 0.0713 | 0.148 |  | 577 | 671 | 0.024 |
| *Richardia brasiliensis* | 0.12631 | -0.99199 | 0.0337 | 0.426 |  | 1344 | 2043 | 0.490 |
| *Ricinus communis* | -0.07481 | -0.9972 | 0.031 | 0.439 |  | 1344 | 1576 | 0.048 |
| *Rotheca sp.1* | -0.71565 | 0.69846 | 0.0875 | 0.136 |  | 526 | 526 | 0.006 |
| *Rubus niveus* | 0.25521 | -0.96689 | 0.0046 | 0.849 |  | 1695 | 1864 | 0.114 |
| *Rubus rosifolius* | 0.3812 | -0.92449 | 0.0013 | 0.951 |  | 895 | 1695 | 0.138 |
| *Rubus steudneri* | 0.9637 | -0.26699 | 0.0993 | 0.121 |  | 2192 | 2528 | 1.567 |
| *Salvia nilotica* | 0.67238 | -0.74021 | 0.0736 | 0.159 |  | 2070 | 2192 | 0.036 |
| *Scutia myrtina* | 0.21089 | 0.97751 | 0.0149 | 0.679 |  | 1462 | 1462 | 0.018 |
| *Searsia natalensis* | 0.05461 | -0.99851 | 0.0521 | 0.277 |  | 1864 | 1864 | 0.018 |
| *Searsia sp.1* | -0.32591 | -0.9454 | 0.0656 | 0.163 |  | 1344 | 1404 | 0.640 |
| *Senecio madagascariensis* | 0.11718 | -0.99311 | 0.0273 | 0.491 |  | 1558 | 2135 | 0.042 |
| *Senecio subsessilis* | 0.8811 | 0.47293 | 0.0423 | 0.381 |  | 2528 | 2528 | 0.024 |
| *Senecio syringifolius* | 0.46827 | 0.88358 | 0.0436 | 0.327 |  | 2414 | 2414 | 0.006 |
| *Senna didymobotrya* | 0.12044 | -0.99272 | 0.0594 | 0.202 |  | 1864 | 2192 | 0.724 |
| *Senna longiracemosa* | -0.78466 | 0.61992 | 0.0778 | 0.132 |  | 577 | 730 | 0.030 |
| *Senna occidentalis* | -0.8479 | -0.53015 | 0.0179 | 0.627 |  | 730 | 1053 | 0.018 |
| *Sesamum angolense* | 0.01959 | 0.99981 | 0.0498 | 0.278 |  | 1528 | 1528 | 0.006 |
| *Sesbania sesban* | -0.99165 | -0.12898 | 0.0129 | 0.699 |  | 529 | 1576 | 0.030 |
| *Sida acuta* | -0.9191 | -0.39403 | 0.0004 | 0.986 |  | 1053 | 1633 | 0.024 |
| *Sida alba* | 0.06519 | 0.99787 | 0.0582 | 0.189 |  | 671 | 1608 | 0.090 |
| *Sida ovata* | -0.83258 | 0.55391 | 0.0222 | 0.521 |  | 529 | 529 | 0.042 |
| *Sida rhombifolia* | 0.21621 | -0.97635 | 0.0117 | 0.74 |  | 1608 | 1608 | 0.006 |
| *Sida tenuicarpa* | 0.31254 | -0.9499 | 0.0066 | 0.812 |  | 1462 | 2043 | 0.132 |
| *Sigesbeckia orientalis* | 0.14927 | -0.9888 | 0.022 | 0.546 |  | 1563 | 1563 | 0.006 |
| *Solanum campylacanthum* | 0.04975 | -0.99876 | 0.0313 | 0.452 |  | 669 | 1864 | 0.347 |
| *Solanum mauritianum* | 0.50905 | -0.86074 | 0.0651 | 0.165 |  | 2070 | 2070 | 0.024 |
| *Solanum nigrum* | 0.09331 | -0.99564 | 0.058 | 0.249 |  | 1344 | 1608 | 0.042 |
| *Solanum renschii* | -0.92254 | 0.38589 | 0.022 | 0.565 |  | 730 | 916 | 0.024 |
| *Sonchus luxurians* | 0.56909 | -0.82227 | 0.0451 | 0.319 |  | 2043 | 2192 | 0.012 |
| *Sonchus oleraceus* | 0.48968 | -0.8719 | 0.0154 | 0.662 |  | 2035 | 2035 | 0.006 |
| *Spermacoce princeae* | 0.62317 | -0.78209 | 0.0492 | 0.287 |  | 1762 | 2192 | 0.227 |
| *Spermacoce pusilla* | -0.05699 | -0.99837 | 0.0277 | 0.516 |  | 1344 | 1344 | 0.006 |
| *Sphaeranthus suaveolens* | 0.50905 | -0.86074 | 0.0651 | 0.165 |  | 2070 | 2070 | 1.423 |
| *Sphagneticola trilobata* | 0.14927 | -0.9888 | 0.022 | 0.546 |  | 1563 | 1563 | 0.012 |
| *Sporobolus pyramidalis* | 0.14927 | -0.9888 | 0.022 | 0.546 |  | 1563 | 1563 | 0.006 |
| *Stachytarpheta urticifolia* | 0.24943 | -0.96839 | 0.0019 | 0.947 |  | 981 | 1630 | 0.090 |
| *Stylosanthes fruticosa* | 0.21319 | -0.97701 | 0.0015 | 0.965 |  | 1624 | 1632 | 0.012 |
| *Syzium sp.1* | 0.11432 | -0.99344 | 0.1113 | 0.117 |  | 1762 | 1762 | 0.120 |
| *Tabernaemontana sp.1* | 0.9074 | -0.42027 | 0.0793 | 0.143 |  | 1695 | 2390 | 0.090 |
| *Tecoma stans* | -0.99714 | 0.07552 | 0.0189 | 0.585 |  | 895 | 1094 | 0.120 |
| *Tephrosia hildebrandtii* | 0.26526 | -0.96418 | 0.0021 | 0.928 |  | 1462 | 1633 | 0.096 |
| *Tephrosia uniflora* | -0.69142 | 0.72246 | 0.0442 | 0.298 |  | 529 | 916 | 0.096 |
| *Tephrosia villosa* | -0.63948 | 0.7688 | 0.0637 | 0.187 |  | 526 | 671 | 0.042 |
| *Thunbergia alata* | 0.49783 | 0.86727 | 0.0001 | 1 |  | 1633 | 1633 | 0.006 |
| *Thylachium thomasii* | -0.31493 | 0.94911 | 0.0344 | 0.406 |  | 669 | 669 | 0.335 |
| *Tinnea aethiopica* | -0.27906 | 0.96027 | 0.0008 | 0.983 |  | 1624 | 1624 | 0.012 |
| *Tithonia diversifolia* | 0.13451 | 0.99091 | 0.06 | 0.188 |  | 1503 | 1700 | 6.727 |
| *Toddalia asiatica* | -0.33829 | -0.94104 | 0.0604 | 0.208 |  | 1352 | 1352 | 0.012 |
| *Tribulus terrestris* | -0.31493 | 0.94911 | 0.0344 | 0.406 |  | 669 | 669 | 0.012 |
| *Trichodesma zeylanica* | 0.14322 | 0.98969 | 0.0731 | 0.149 |  | 1462 | 1539 | 0.999 |
| *Tridax procumbens* | -0.55341 | 0.83291 | 0.0218 | 0.581 |  | 529 | 1632 | 0.305 |
| *Trifolium semipilosum* | 0.98516 | -0.17163 | 0.1128 | 0.089 | . | 2390 | 2528 | 0.036 |
| *Triumfetta brachyceras* | 0.5387 | -0.8425 | 0.0875 | 0.123 |  | 1563 | 2192 | 0.377 |
| *Triumfetta rhomboidea* | 0.27299 | -0.96202 | 0.051 | 0.27 |  | 1218 | 2135 | 0.078 |
| *Triumfetta sp.1* | -0.05699 | -0.99837 | 0.0277 | 0.516 |  | 1344 | 1344 | 0.054 |
| *Turraea robusta* | -0.80201 | -0.59731 | 0.1147 | 0.077 | . | 1094 | 1094 | 0.006 |
| *Urochloa Trichopus* | -0.83258 | 0.55391 | 0.0222 | 0.521 |  | 529 | 529 | 0.006 |
| *Vernonia adoensis* | 0.39289 | -0.91959 | 0.002 | 0.945 |  | 1695 | 1695 | 0.012 |
| *Vernonia auriculifera* | 0.05461 | -0.99851 | 0.0521 | 0.277 |  | 1864 | 1864 | 0.006 |
| *Vernonia brachycalyx* | 0.08329 | -0.99653 | 0.0668 | 0.161 |  | 1462 | 1864 | 0.921 |
| *Vernonia cinerascens* | -0.99953 | 0.03054 | 0.0376 | 0.379 |  | 627 | 627 | 0.024 |
| *Vernonia glabra* | 0.51957 | 0.85443 | 0.0118 | 0.715 |  | 1530 | 1700 | 0.233 |
| *Vernonia karaguensis* | 0.99945 | -0.03308 | 0.0039 | 0.895 |  | 1462 | 1563 | 0.054 |
| *Vernonia lasiopus* | 0.19732 | 0.98034 | 0.0051 | 0.863 |  | 1352 | 1831 | 0.395 |
| *Vernonia sp.1* | 0.99822 | -0.05967 | 0.0035 | 0.871 |  | 1671 | 1671 | 0.006 |
| *Vernonia usambarensis* | -0.43892 | -0.89852 | 0.089 | 0.118 |  | 730 | 1352 | 0.084 |
| *Veronica abyssinica* | 0.8811 | 0.47293 | 0.0423 | 0.381 |  | 2528 | 2528 | 0.006 |
| *Wahlenbergia abyssinica* | 0.23556 | -0.97186 | 0.0068 | 0.802 |  | 1632 | 1632 | 0.006 |
| *Waltheria indica* | -0.96166 | -0.27424 | 0.0024 | 0.952 |  | 730 | 1563 | 0.383 |
| *Zehneria scabra* | 0.50905 | -0.86074 | 0.0651 | 0.165 |  | 2070 | 2070 | 0.257 |
| *Zornia setosa* | 0.23556 | -0.97186 | 0.0068 | 0.802 |  | 1632 | 1632 | 0.006 |
